# The fish pathogen *Aliivibrio salmonicida* LFI1238 can degrade and metabolize chitin despite major gene loss in the chitinolytic pathway

**DOI:** 10.1101/2021.03.24.436902

**Authors:** Anna Skåne, Giusi Minniti, Jennifer S.M. Loose, Sophanit Mekasha, Bastien Bissaro, Geir Mathiesen, Magnus Ø. Arntzen, Gustav Vaaje-Kolstad

## Abstract

The fish pathogen *Aliivibrio (Vibrio) salmonicida* LFI1238 is thought to be incapable of utilizing chitin as a nutrient source since approximately half of the genes representing the chitinolytic pathway are disrupted by insertion sequences. In the present study, we combined a broad set of analytical methods to investigate this hypothesis. Cultivation studies revealed that *Al. salmonicida* grew efficiently on *N*-acetylglucosamine (GlcNAc) and chitobiose ((GlcNAc)_2_), the primary soluble products resulting from enzymatic chitin hydrolysis. The bacterium was also able to grow on chitin particles, albeit at a lower rate compared to the soluble substrates. The genome of the bacterium contains five disrupted chitinase genes (pseudogenes) and three intact genes encoding a glycoside hydrolase family 18 (GH18) chitinase and two auxiliary activity family 10 (AA10) lytic polysaccharide monooxygenases (LPMOs). Biochemical characterization showed that the chitinase and LPMOs were able to depolymerize both α- and β-chitin to (GlcNAc)_2_ and oxidized chitooligosaccharides, respectively. Notably, the chitinase displayed up to 50-fold lower activity compared to other well-studied chitinases. Deletion of the genes encoding the intact chitinolytic enzymes showed that the chitinase was important for growth on β-chitin, whereas the LPMO gene-deletion variants only showed minor growth defects on this substrate. Finally, proteomic analysis of *Al. salmonicida* LFI1238 growth on β-chitin showed expression of all three chitinolytic enzymes, and intriguingly also three of the disrupted chitinases. In conclusion, our results show that *Al. salmonicida* LFI1238 can utilize chitin as a nutrient source and that the GH18 chitinase and the two LPMOs are needed for this ability.

**IMPORTANCE:** The ability to utilize chitin as a source of nutrients is important for the survival and spread of marine microbial pathogens in the environment. One such pathogen is *Aliivibrio (Vibrio) salmonicida*, the causative agent of cold water vibriosis. Due to extensive gene decay, many key enzymes in the chitinolytic pathway have been disrupted, putatively rendering this bacterium incapable of chitin degradation and utilization. In the present study we demonstrate that *Al. salmonicida* can degrade and metabolize chitin, the most abundant biopolymer in the ocean. Our findings shed new light on the environmental adaption of this fish pathogen.

## INTRODUCTION

Chitin is one of the most abundant biopolymers in nature and is a primary component of rigid structures such as the exoskeleton of insects and crustaceans, and the cell wall of fungi and some algae (1–4). Some reports also indicate that chitin is found in the scales and gut of fish (5, 6). This linear polysaccharide consists of *N*-acetyl-D-glucosamine (GlcNAc) units linked by β-1,4 glycosidic bonds that associates with other chitin chains to form insoluble chitin fibers. Despite the recalcitrance of chitin, the polymer is readily degraded and metabolized by chitinolytic microorganisms in the environment (7, 8).

Most bacteria solubilize and depolymerize chitin by secreting chitinolytic enzymes. Such enzymes include chitinases from family 18 and 19 of the glycoside hydrolases (GH18 and - 19) and lytic polysaccharide monooxygenases (LPMOs) from family 10 of the auxiliary activities (AA10), according to classification by the carbohydrate active enzyme database (CAZy; http://www.cazy.org/) (9). Whereas chitinases cleave chitin chains by a hydrolytic mechanism (10, 11), LPMOs perform chitin depolymerization by an oxidative reaction (12–14). The latter enzymes usually target the crystalline parts of chitin fibers that are inaccessible for the chitinases. When combined, chitinases and LPMOs act synergistically, providing efficient depolymerization of this recalcitrant carbohydrate (12, 15–17). The products of enzymatic chitin degradation are mainly GlcNAc and (GlcNAc)_2_, but also native and oxidized chitooligosaccharides, the latter (aldonic acids) arising from LPMO activity.

The chitin degradation pathway is conserved in the *Vibrionaceae* (18, 19). Here, GlcNAc and (GlcNAc)_2_ are transported into the periplasm by unspecific porins (20, 21) or by dedicated transport proteins for chitooligosaccharides ((GlcNAc)_2-6_), named chitoporins (22, 23). Once transported to the periplasm, (GlcNAc)_2-6_ may be hydrolyzed to GlcNAc by family GH20 *N*-acetylhexosaminidases or *N,N*-diacetylchitobiose phosphorylases (24). Transport of GlcNAc or deacetylated GlcN across the inner membrane can occur through phosphotransferase systems, while (GlcNAc)_2_ may be transported through the action of an ABC transporter (18). Once located in the cytosol GlcNAc, GlcNAc1P or GlcN enter the amino-sugar metabolism. It should be noted that the fate of chitooligosaccharide aldonic acid is not known.

Chitin degradation can be achieved by several marine bacteria, and can give advantages for survival and proliferation in the marine environment (8, 25). Some pathogens have chitin central in their lifecycle, the most prominent example being the human pathogen *Vibrio cholerae* that uses chitin-containing zoo-plankton as transfer vectors and nutrition (26, 27). The ability of the Gram-negative marine bacterium *Aliivibrio salmonicida* (previously *Vibrio salmonicida*), to utilize chitin or GlcNAc as a nutrient source is controversial. This pathogenic bacterium, which is the causative agent of cold water vibriosis in salmonids, was identified as a new vibrio-like bacteria in 1986 (28). Upon discovery and initial characterization of the pathogen (strain HI 7751), Egidius et al. did not observe degradation of chitin by the bacterium when growing on agar plates containing purified chitin. On the other hand, the monomeric building block of chitin, GlcNAc, was readily consumed by the bacterium. When the genome of the bacterium was sequenced two decades later (strain LFI1238), it was shown that insertion sequence (IS) elements caused disruption of almost 10% of the protein encoding genes (29, 30). Especially effected was the chitin utilization pathway where seven genes, including three chitinases and a chitoporin, were either disrupted or truncated (29). In addition, the gene encoding the periplasmic chitin-binding protein (VSAL_I2576, also called CBP) was disrupted by a frameshift. The CBP ortholog in *V. cholerae* (VC_0620) has been shown to activate the two-component chitin catabolic sensor/kinase ChiS that regulates chitin utilization (31, 32). The gene encoding the ChiS ortholog in *Al. salmonicida* is intact (29), along with the Tfox encoding gene which protein product also is involved in regulation of enzymes related to chitin degradation in the *Vibrionaceae* (33, 34). Of the putative secreted chitinolytic enzymes, only one chitinase and two lytic polysaccharide monooxygenases remained intact in the *Al. salmonicida* genome. It was suggested that such extensive gene disruption could indicate inactivation of this pathway and indeed, the authors could not observe neither degradation of insoluble chitin nor utilization of GlcNAc as a nutrient source (29).

In order to obtain a deeper understanding of the roles of the *Al. salmonicida* chitinolytic enzymes, we have analyzed the chitin degradation potential of *Al. salmonicida* LFI1238 by biochemical characterization of the secreted chitinolytic enzymes, gene deletion and cultivation experiments, gene expression analysis and proteomics.

## RESULTS

### *Al. salmonicida* can utilize both GlcNAc and (GlcNAc)_2_ as nutrient sources

To assess the ability of *Al. salmonicida* LFI1238 (abbreviated *“Al. salmonicida”* to avoid confusion with *Aeromonas salmonicida*) to grow on GlcNAc and (GlcNAc)_2_, the wild type strain was cultivated in minimal medium supplemented with 0.2% glucose (11.1 mM; control experiment), 0.2 % GlcNAc (9.0 mM), or 0.2 % (GlcNAc)_2_ (4.7 mM) over a period of 92 hours. The cultivation experiments showed that *Al. salmonicida* can utilize both GlcNAc and (GlcNAc)_2_ as sole carbon sources (Fig. 1). Growth rates were compared by calculating the specific rate constants (µ) and generation time across the exponential phase (Table S1), showing little difference between the three carbon sources. In order to correlate GlcNAc and (GlcNAc)_2_ consumption with the bacterial growth, the concentration of these sugars in the culture supernatant were determined at different time points during growth (Fig. 1E, F). The data show decreasing concentrations of GlcNAc during growth and complete depletion within 40 hours (Fig. 1E). In comparison, (GlcNAc)_2_ is utilized at a slower speed, becoming depleted after 80 hours (Fig. 1F).

**Figure 1.**
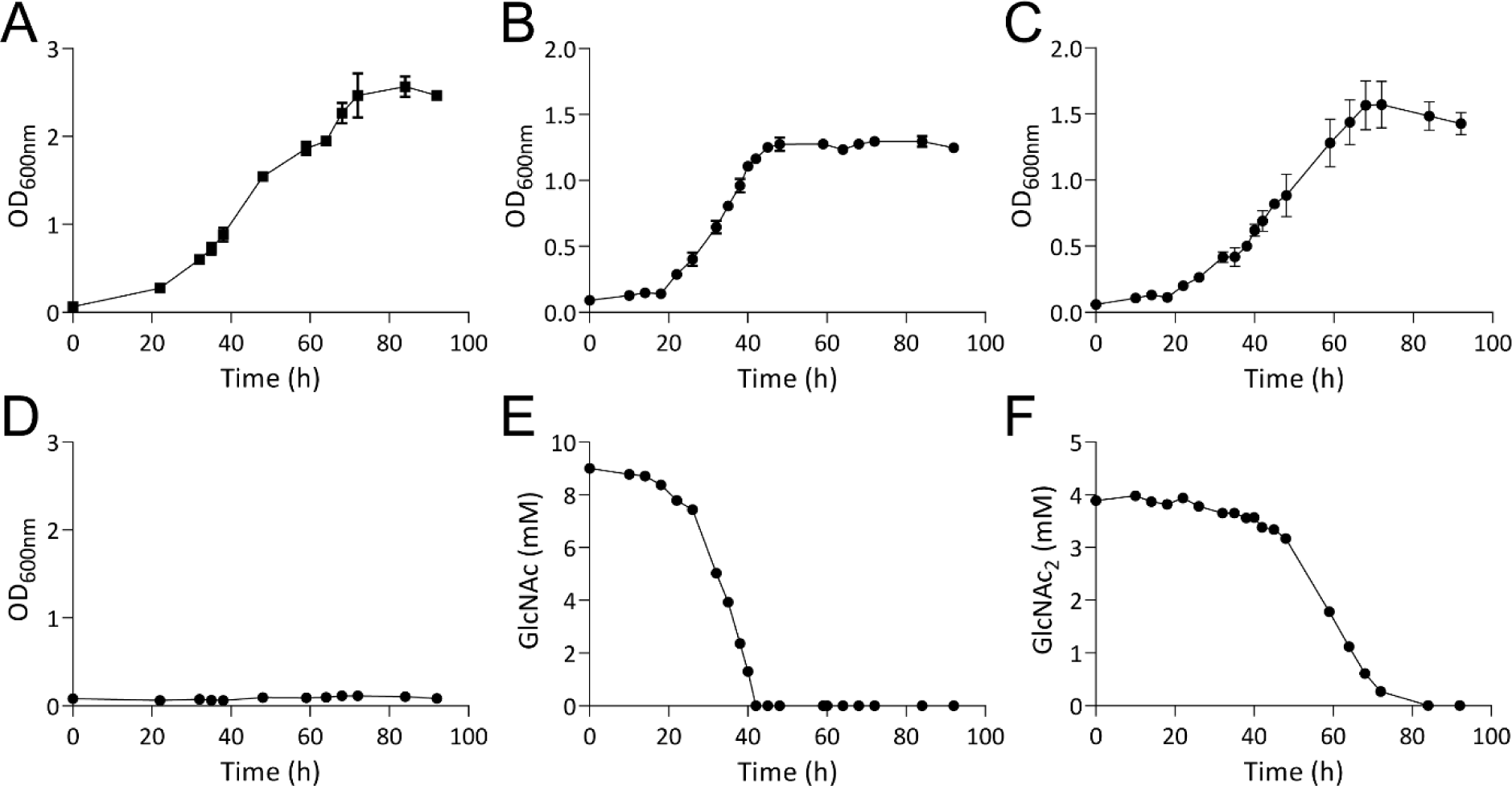
Utilization of Glucose, GlcNAc and (GlcNAc)_2_. Panels A to C show the growth of *Al. salmonicida* LFI1238 in minimal media supplemented with 0.2 % glucose, 0.2% GlcNAc (9.0 mM) or 0.2 % (GlcNAc)_2_ (4.7 mM), respectively. Growth in defined media without supplementation of carbon source (negative control) is shown in panel D. Growth results are shown as mean value of three biological replicates and the standard deviation is indicated. Panels E & F show the depletion of soluble substrates by *Al. salmonicida*, determined by sampling of the culture supernatant of one replicate different time points through the growth time-period and quantification of GlcNAc (panel E) or (GlcNAc)_2_ (panel F) by ion exclusion chromatography. Results are shown as the mean value of three technical replicates.

### Sequence analysis and homology modelling

Since *Al. salmonicida* was able to utilize both GlcNAc and (GlcNAc)_2_, the major products of enzymatic chitin degradation, it was of interest to analyze the chitinolytic potential of the bacterial genome, investigating the details of both intact genes and pseudogenes. A previous study had already identified the presence of three putatively secreted chitinolytic enzymes (29). Annotation of putative CAZy domains of these three enzymes using the dbCAN server (35) showed that the chitinase sequence, here named *As*Chi18A, (that contains 881 amino acids, which is unusually large for a chitinase) contains predicted CBM5 and CBM73 chitin binding domains and a C-terminal GH18 domain, the latter modest in size (only 324 amino acids; Fig. 2A). The protein sequence also shows long regions that were not annotated. Attempts to functionally annotate these regions with other sequence analysis servers such as InterPro, Pfam and SMART were inconclusive. The relatively small size of the GH18 catalytic domain indicates an enzyme stripped of most sub-domains that often are in place to form a substrate binding cleft. Indeed, homology modelling using Swiss-Model (36) revealed a model structure with a shallow substrate binding cleft, reminiscent of a non-processive *endo*-chitinase, which is clearly observed when compared to the processive *exo*-chitinase *Sm*Chi18A from *Serratia marcescens* that has a deep substrate binding cleft and the shallow-clefted, non-processive chitinase ChiNCTU2 from *Bacillus cereus* ((37); Fig. 2B). *As*Chi18A also shows an arrangement of active site residues that is similar to that of the latter enzyme (Fig. S1).

**Figure 2.**
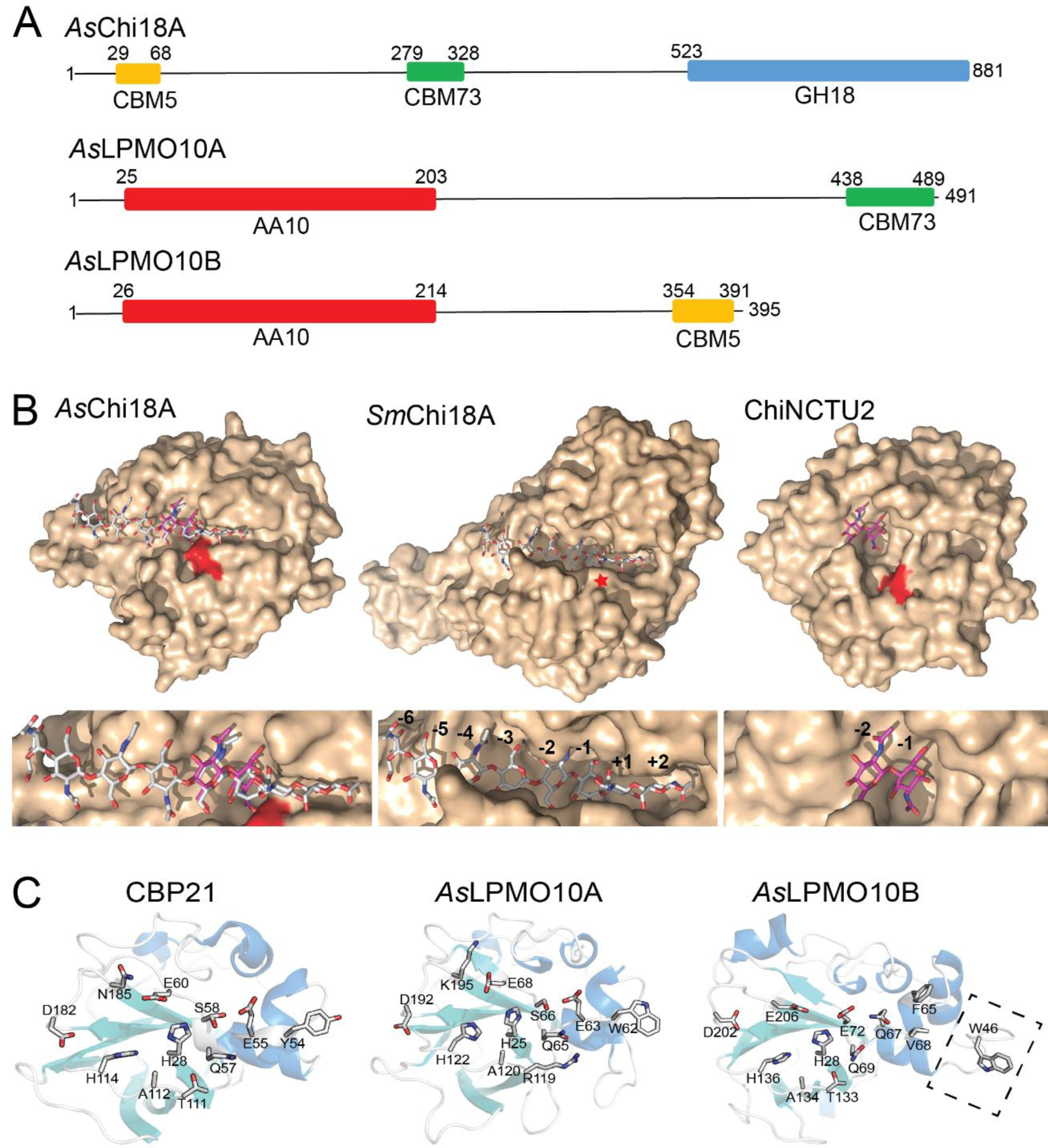
Predicted domains and three dimentional structures of the *A. salmonicida* chitinase and LPMOs. **(A)** Prediction of CAZy domains of the chitinolytic enzymes was performed using the dbCAN server. Numbers indicate the position in the sequence. The theoretical molecular weight of the proteins calculated by the ProtParam tool (in the absence of the predicted signal peptide) is 87.4, 52.5 and 41.2 kDa for *As*Chi18A, *As*LPMO10A and *As*LPMO10B, respectively. Signal peptides were determined by the SignalP 4.0 server (http://www.cbs.dtu.dk/services/SignalP/) and represent residues 1-29, 1-25 and 1-26 for *As*Chi18A, *As*LPMO10A and *As*LPMO10B, respectively. The GenBank protein identifiers for the enzymes are C442.1 (*As*Chi18A, also called “endochitinase A”), C888.1 (*As*LPMO10A, also called “chitin binding protein”) and C971.1 (*As*LPMO10B, also called “chitinase B”). **(B)** The homology model of *As*Chi18A (left structure) and the structures of *Sm*Chi18A deep clefted *exo*-chitinase from *S. marcescens* (middle structure) and the *Bacillus cereus* GH18 ChiNCTU2 shallow clefted chitinase (37) are shown in light brown surface representation with the catalytic acids colored red (or indicated by a red star for *Sm*ChiA, as it is concealed by other amino acids). Ligands are shown in stick representation with gray (chitooctaose; *Sm*Chi18A) and purple (chitobiose; ChiNCTU2) colored carbon atoms. Subsites are indicated by numbering. Ligands shown in the *As*Chi18A substrate binding cleft are derived from structural superimpositions of the *As*Chi18A model with *Sm*Chi18A or ChiNCTU2 and are provided for illustrational purposes only. The template used for modelling the *As*Chi18A catalytic GH18 domain was PDB ID 3N1A (apo-enzyme structure of ChiNCTU2 from *B. cereus*) and gave a Qmean value of −1.99, which represents a good quality model. **(C)** The crystal structure of CBP21 (PDB ID 2BEM) and the homology models for *As*LPMO10A and *As*LPMO10B are shown in cartoon representation. For CBP21, the side chains of the amino acids that have been shown to be invovled in substrate binding by experimental evidence (42, 43, 103) are shown in stick representation. The corresponding amino acids in *As*LPMO10A and *As*LPMO10B are also shown in stick representation. One exception is W46 of *As*LPMO10B, which is not present in the two other enzymes. The latter residue is positioned on an insertion that potentially extends the putative binding surface (indicated by rectangle with dashed lines). In CBP21, Ser58 is shown with two alternative side chain conformations. Swiss Model was used with default paramters to generate the homology models of *As*LPMO10A and -B, using PDB structures 2XWX (66.5% sequence identity to *As*LPMO10A) and 4YN2 (43.6% sequence identity to *As*LPMO10B) as templates, respectively. The Q-mean scores obtained were −1.65 for *As*LPMO10A model and -3.34 for *As*LPMO10B

Annotation of the LPMO sequences showed that both proteins contained an N-terminal catalytic AA10 domain and a C-terminal CBM73 or CBM5 chitin-binding domain in *As*LPMO10A and –B, respectively (Fig. 2A). Like the chitinase, both LPMOs displayed regions in the sequence that were not possible to annotate using standard bioinformatics tools. Pair-wise sequence alignment of the two LPMOs revealed only a 20% identity between the catalytic domains. Blast search and modelling by homology of the individual catalytic domains showed that the catalytic module of *As*LPMO10A was similar to CBP21 from *S. marcescens* (49.5% identity, Fig. 2 C; (38, 39)) and to the catalytic AA10 domain of GbpA, a *Vibrio cholerae* colonization factor ((40); 65.6% identity). The similarity of full-length *As*LPMO10A to *V. cholera* GbpA (61% sequence identity) and their similar multi-modular architecture (both have a N-terminal AA10 LPMO domain, followed by a “GbpA2” domain, an un-annotated domain and a C-terminal CBM73 domain) indicate the possibility of functionally similar roles. The catalytic AA10 domain of *As*LPMO10B is, as already noted, very unlike *As*LPMO10A. From sequence database searches, orthologs were identified in a large variety of species from the *Vibrionaceae* family, and also in other marine bacteria like *Shewanella* and *Pseudoalteromonas*. None of these related enzymes have hitherto been biochemically characterized. When searching for similar sequences in the PDB database, the most similar structure to the *As*LPMO10B catalytic domain belongs to the viral proteins called “spindolins” (43.5% identity, but the alignment contains many insertions/ deletions). There exist no activity data for spindolins, but it is assumed that they are active towards chitin (41). It is therefore not straightforward to assign an activity to *As*LPMO10B based on sequence analysis. In order to analyze the putative structural difference between the LPMO domains, homology models were made using the Swiss-Model homology modelling software (36). When compared to CBP21, one of the best characterized family AA10 LPMOs, both *Al. salmonicida* LPMOs show several differences that may influence both substrate binding and catalysis (Fig. 2C): *As*LPMO10A is relatively similar to CBP21 but displays some differences that may be of functional relevance: amino acids W62, R119, K195 in *As*LPMO10A correspond to amino acids Y54, T111 and N185 in CBP21 that all have been shown to have influence on substrate binding and the functional stability of the enzyme (42, 43). *As*LPMO10B shows an active site environment similar to CBP21 but has an extension of the putative binding surface that positions a putatively solvent exposed Trp (W46) further away from the active site histidines than for Y54 in CBP21 and W62 in *As*LPMO10A. Whether these differences are important for the substrate binding properties of the enzymes is not straightforward to interpret based on the data presented in this study, since both *Al. salmonicida* proteins have CBMs that very likely contribute to chitin binding.

### Analysis of pseudogenes related to chitin catabolism

In addition to the intact genes encoding the chitinase, *As*Chi18A and LPMOs, *As*LPMO10A and -B, the genome of *Al. salmonicida* LFI12338 harbors multiple pseudogenes encoding truncated or fragmented enzymes related to chitin catabolism that are assumed to be non-functional (ORF identifiers VSAL_I2352, VSAL_I0763, VSAL_I0902, VSAL_I1108, VSAL_I1414 and VSAL_I1942). Interestingly, transcription of *Al. salmonicida* pseudogenes (including chitinase-related pseudogenes) has been observed (44–46). In addition, *Al. salmonicida* is motile despite two flagellar synthesis genes (*fliF*/VSAL_I2308 and (*flaG*/VSAL_I12316) being disrupted by premature stop codons (29). Thus, we performed a deeper analysis of the *Al. salmonicida* pseudogenes related to the chitinolytic machinery to investigate their putative functionality. The analysis revealed that VSAL_I2352 (predicted chitoporin) contains a frameshift after codon 266, which most likely will result in a non-functional protein if expressed. On the other hand, VSAL_I0763 (chitinase fragment), VSAL_I0902 (truncated chitinase), VSAL_I1108 (truncated chitodextrinase), VSAL_I1414 (disrupted chitinase) and VSAL_I1942 (disrupted chitinase) are rather truncated or disrupted by the type Vsa_2 insertion sequence (IS) elements (Fig. 3A), resulting in coding sequences (CDSs) of varying lengths that may give functional protein if expressed (Fig. 3B). Annotation of putative CAZy domains predicted that VSAL_I0902 (truncated chitinase fragment), VSAL_I1108 (truncated chitodextrinase) and VSAL_I1942 (disrupted chitinase) contain regions encoding GH18 domains, while VSAL_I1414 (disrupted chitinase) was predicted to contain a region encoding a GH19 domain (Fig. 3B). No functional domain was predicted for VSAL_I0763 (sequence containing 609 nucleotides truncated by upstream IS element and subsequent recombinations). It is believed that VSAL_I0902 and VSAL_I0763 are fragments belonging to one single chitinase (29).

**Figure 3.**
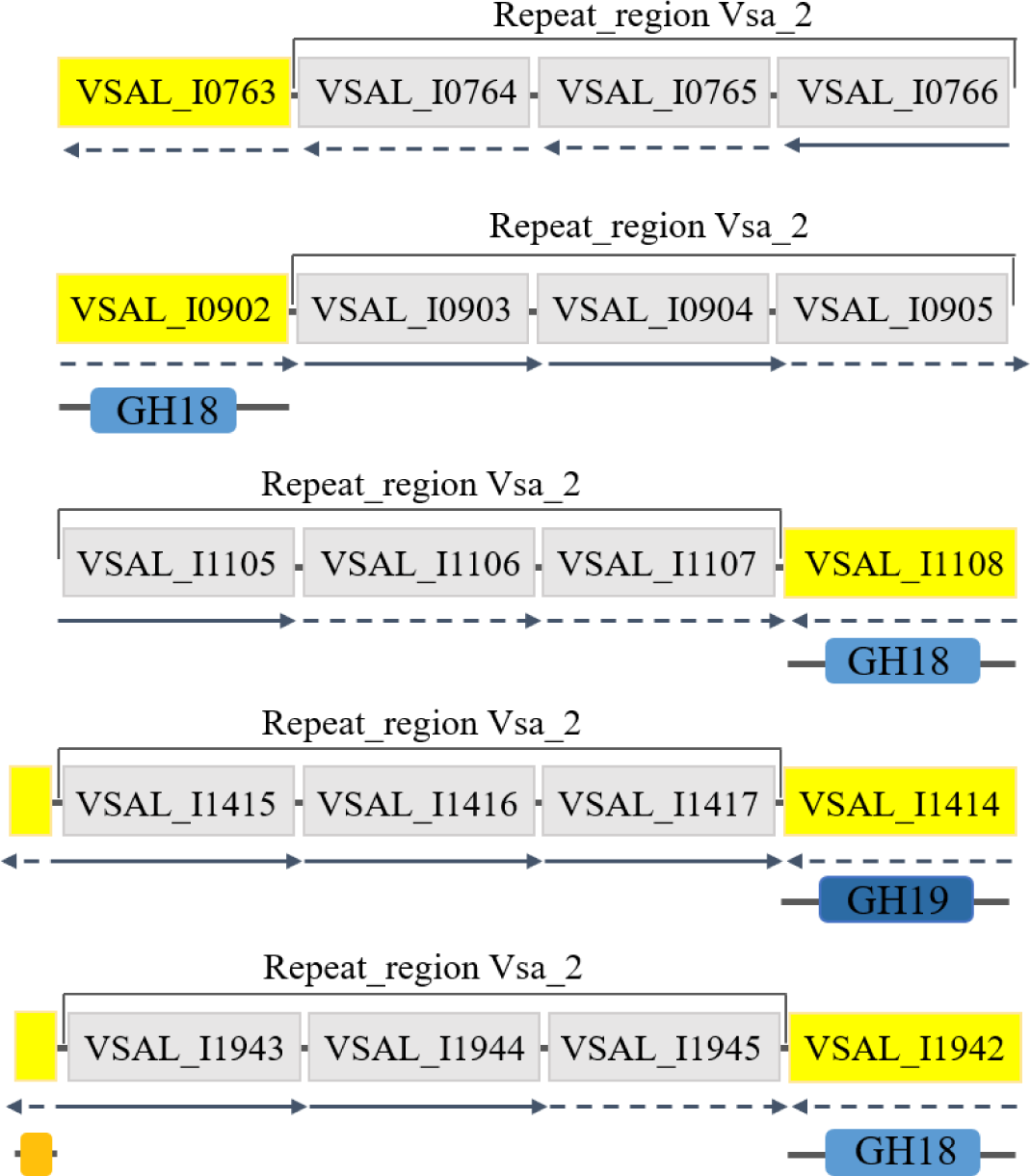
Sequence analysis of putative chitinase pseudogenes. The gene locus and insertion sequence elements are shown in yellow and gray, respectively, with the locus name indicated. Solid lined arrow direction indicates reading frame direction, while dashed lined arrows indicate pseudogenes. CAZyme annotation of the pseudogenes genes was done using dbCAN2 and the resulting enzyme activity prediction is displayed below each gene. Annotation of VSAL_I0763, VSAL_I0902, VSAL_I1108 was performed with the truncated chitinase/chitodextrinase sequence. VSAL_I1414 and VSAL_I1942 were analyzed using the full-length sequences including repeat region. The illustrations representing the ORFs are not to scale.

In conclusion, four truncated chitinase genes contain regions encoding GH-domains which may give functional protein if translated.

### *As*Chi18A and *As*LPMO10A and -B binds chitin

In order to determine the biochemical properties of putatively chitinolytic enzymes (the pseudogene encoded chitinases were not expressed and characterized), *As*Chi18A and *As*LPMO10A and -B were cloned, expressed and purified (Fig. S2). The presence of putative chitin binding modules on all three chitinolytic enzymes prompted investigation of the substrate binding properties of the proteins. Using purified protein, α-chitin and β-chitin were used as substrates in particle sedimentation assays (Fig. 4). All proteins showed binding to the substrate particles and *As*LPMO10B seems to bind slightly weaker to the substrates used compared to *As*LPMO10A.

**Figure 4.**
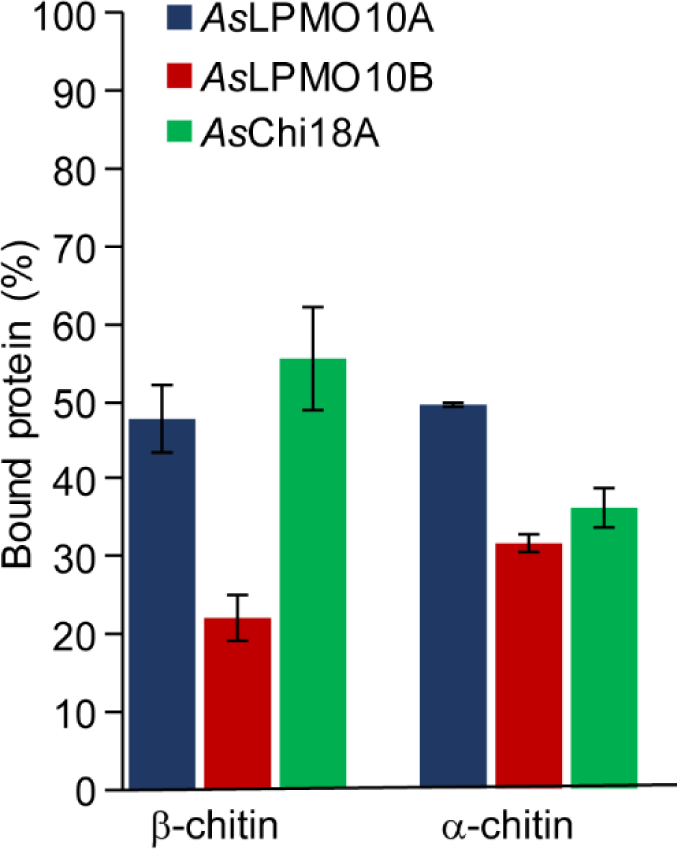
Substrate binding of *As*Chi18A, *As*LPMO10A and -B. Each bar shows the percentage of bound proteins after 2 h of incubation at 30 °C. Reactions contained 10 mg/mL of substrate, 0.75 µM (LPMOs) or 0.50 µM (*As*Chi18A) of enzymes and 10 mM of Tris-HCl buffer at pH 7.5. All reactions were run in triplicates and the standard deviations are indicated by error bars.

### *As*Chi18A displays low chitinolytic activity

Since all three enzymes bound to chitin, the catalytic properties of the purified chitinase and two LPMOs were analyzed. Using β-chitin as substrate, the activity and operational stability of *As*Chi18A was followed over several hours at temperatures ranging from 10-60 °C. The progress curves observed for *As*Chi18A indicate an optimal operational stability, i.e. the highest temperature for which enzyme activity remains stable over time, at approximately 30 °C (Fig. 5A). Similar to other GH18 chitinases, the dominant product of chitin hydrolysis by *As*Chi18A was (GlcNAc)_2_ with small amounts of GlcNAc (< 5%).

**Figure 5.**
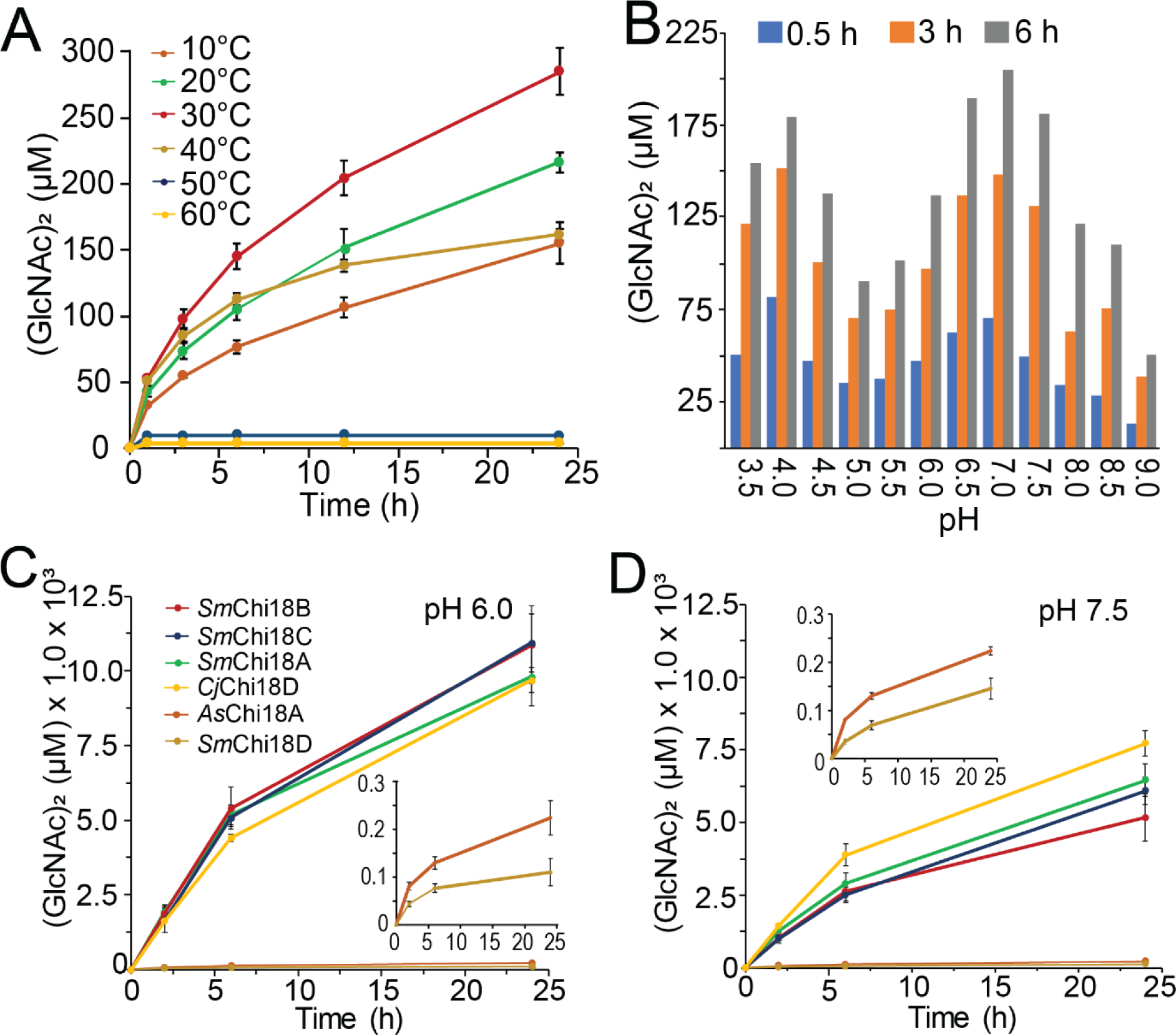
Enzymatic properties of *As*Chi18A. Production of (GlcNAc)_2_ by *As*Chi18A analysed at various temperatures (A) and pH values (B). The activity of *As*Chi18A was also compared to the chitinases from *Serratia marcescens* (*Sm*Chi18A, -B, -C and -D) and *C. japonicus* (*Cj*Chi18D) at pH 6.0 (C) and 7.5 (D). All reactions conditions included 10 mg/mL β-chitin and 0.5 µM enzyme. For data displayed in panel A, reactions were carried out at pH 7.5. For the data displayed in panel B, all reactions were incubated at 30 °C. Buffers used were formic acid pH 3.5, acetic acid pH 4.0 and 4.5, ammonium acetate pH 4.5 and 5.0, MES pH 5.5, 6.0 and 6.5, BisTris-HCl pH 7.0, Tris-HCl pH 7.5 and 8.0 and Bicine pH 8.5 and 9.0. The amounts of (GlcNAc)_2_ presented are based on the average of three independent reactions containing 10 mg/mL β-chitin, 0.5 µM enzyme and 10 mM buffer. The insets in panel C and D show magnified views of reactions catalysed by *As*Chi18A and *Sm*Chi18D. Standard deviations are indicated by error bars (n=3).

In order to compare *As*Chi18A activity with other well-characterized chitinases, the chitin degradation potential of the enzyme was compared with the four GH18 chitinases of *S. marcescens* (*Sm*Chi18A, -B, -C and -D) (47–49) and, *Cj*Chi18D, which is the most potent chitinase of *Cellvibrio japonicus* (50). Activities were monitored at pH 6.0 (Fig. 5C), which is the pH where the *S. marcescens* and *C. japonicus* chitinases have their optima (47, 51, 52), and at pH 7.5 (Fig. 5D), which is a typical pH of sea water and the near pH-optimum of *As*Chi18A. Strikingly, *Sm*Chi18A, -B, -C and *Cj*Chi18D yielded more than 50-fold more (GlcNAc)_2_ than *As*Chi18A after 24 h incubation at pH 6. At pH 7.5, the differences in yields were lower (in the range of 25-40-fold larger yields, except for *Sm*Chi18D), most likely reflecting the difference in pH optima. It should be noted that the presence of NaCl in concentrations similar to sea water (∼0.6 M) only marginally influenced *As*Chi18A activity (Fig. S3).

### *As*LPMO10A and -B are active towards chitin

Both *Al. salmonicida* LPMOs were able to oxidize α- and β-chitin, yielding aldonic acid chitooligosaccharide products with degree of polymerization ranging from 3 to 8 (Fig. S4). Such product profiles are commonly observed for family AA10 LPMOs that target chitin (12, 14, 53). The two enzymes displayed slightly different operational stabilities when probed at temperatures ranging from 10 to 60 °C (Fig. 6). *As*LPMO10A showed an operational stability similar to that of *As*Chi18A, being approximately 30 °C (Fig. 6A, B). In contrast, *As*LPMO10B showed an operational stability lower than 30 °C (Fig. 6C, D). Comparison of the LPMO activities showed that *As*LPMO10A seems generally more active than *As*LPMO10B, the former enzyme yielding approximately twice as much soluble oxidized products than the latter (Fig. 6B, D).

**Figure 6.**
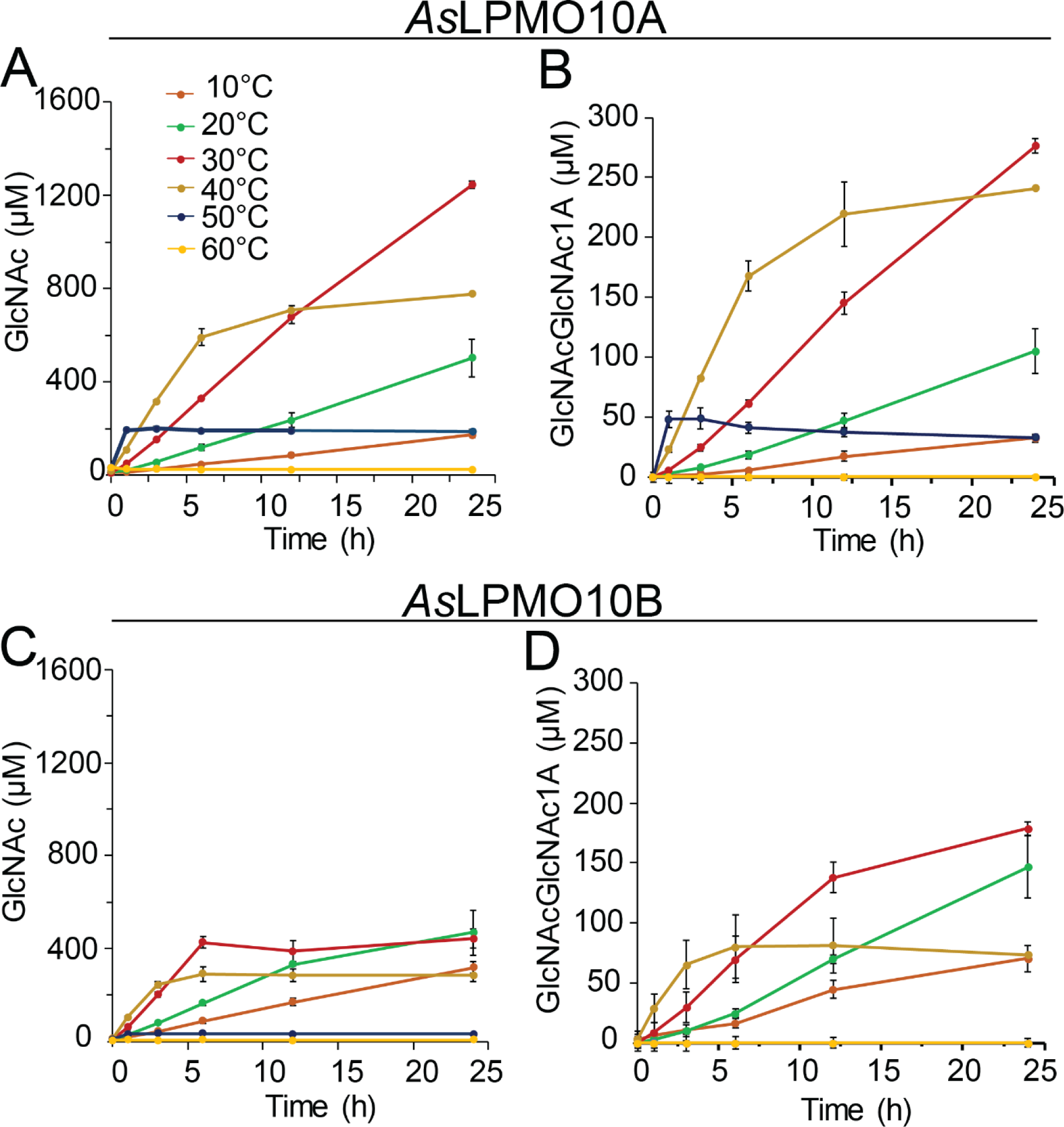
Operational temperature stability of *A. salmonicida* LPMOs. The activity of *As*LPMO10A and *As*LPMO10B is indicated by the production of GlcNAc is shown in panel A and C, respectively. Since the end-product of chitin degradation by the LPMOs are oxidized chitooligosaccharides (Fig. S4) that are inconvenient to quantify, the reaction products obtained from the reactions were depolymerized by Chitobiase that completely converts the oligosaccharide mixture to GlcNAc and oxidized (GlcNAc)_2_ (i.e. GlcNAcGlcNAc1A). The quantities of the latter products formed by the LPMOs, are shown in panel B and D. The amounts presented are based on the average of three independent reactions, which contained 10 mg/mL of β-chitin, 1 µM of enzyme, 1 mM of ascorbic acid and 10 mM of Tris-HCl buffer at pH 7.5, incubated at different temperatures between 10 and 60 °C (colour code provided in panel A). Standard deviations are indicated by error bars.

### Combination of the chitinase and LPMOs shows enzyme synergies

For the putative chitinolytic system of *Al. salmonicida* the situation was different than any other chitinolytic system studied since the chitin degradation potential of the chitinase was substantially lower than that of the LPMOs (Fig. 5C, D and Fig. 6). Usually, the chitinase of a chitinolytic system is substantially more efficient in substrate solubilization than the LPMO. Nevertheless, synergies were observed when combining the *As*Chi18A with *As*LPMO10B giving an almost double yield than the sum of products calculated by adding the sum of their individual yields, for both β- and α-chitin (Fig. 7). *As*LPMO10A, on the other hand, showed a weaker synergy when combined with *As*Chi18A.

**Figure 7.**
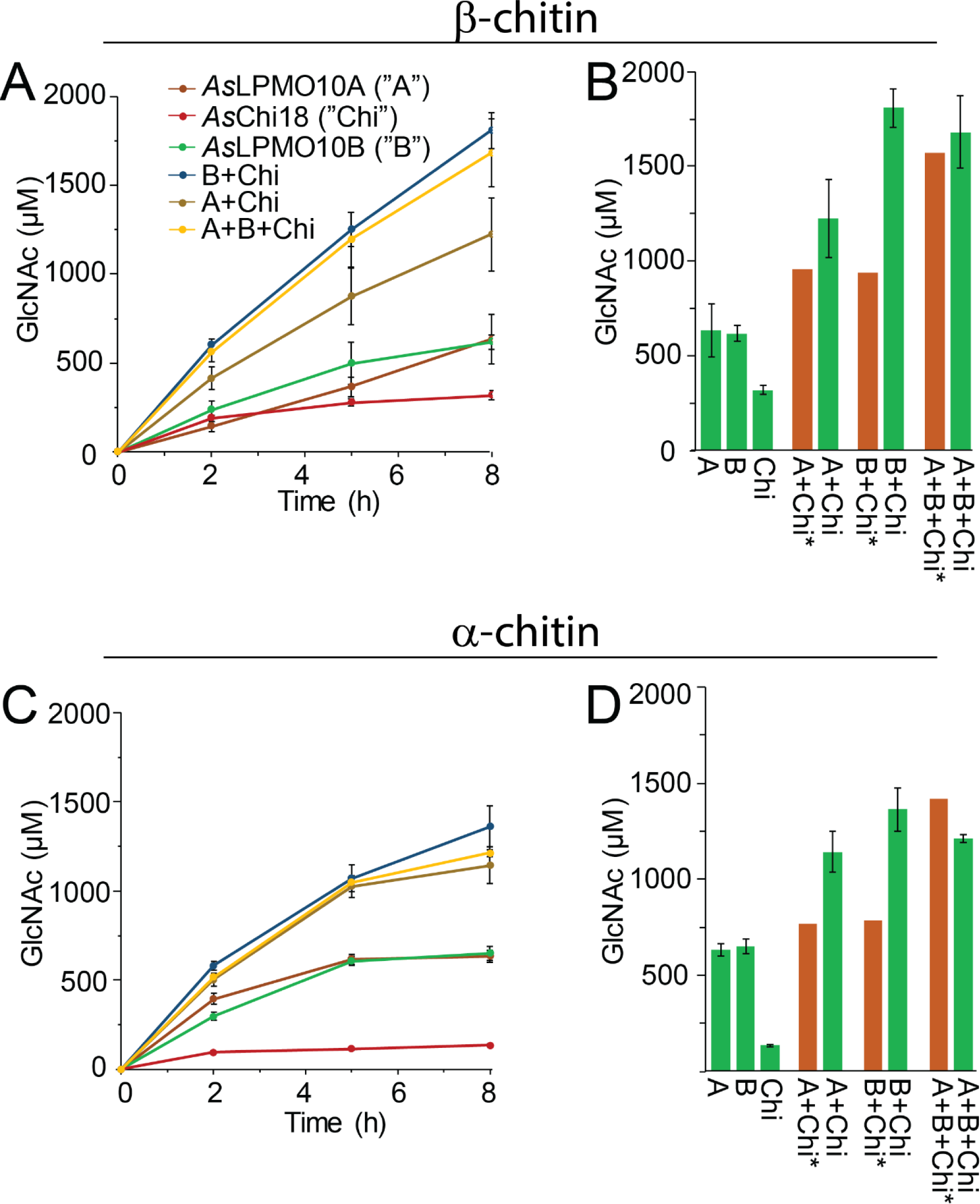
Synergistic activity of *As*LPMO10s and *As*Chi18A on chitin. Panels A and C show the production of GlcNAc by the individual and combined enzymes on β- and α-chitin, respectively. Panels B and D show the theoretically calculated amounts of GlcNAc based on the sum of its production by the individual enzymes (*, brown bars) and the detected amounts of GlcNAc by combining the enzymes after 8 h (green bars). The amounts presented are based on the average of three independent reactions containing 10 mg/mL of chitin substrate, 1 µM of LPMOs and/or 0.5 µM of GH18, 1 mM of ascorbic acid and 10 mM of Tris-HCl buffer at pH 7.5, incubated at 30 °C for 8 h. Standard deviations are indicated by error bars (n=3).

### *As*Chi18A is important for growth of *Al. salmonicida* on chitin

Since the *Al. salmonicida* chitinase and LPMOs were able to depolymerize both α- and β-chitin to soluble sugars that are metabolizable for the bacterium (GlcNAc and (GlcNAc)_2_), the ability of the bacterium to utilize chitin particles as a carbon source was assessed. For this experiment, β-chitin was used for its higher purity and lower recalcitrance compared to α-chitin. To unravel the roles of *As*Chi18A and *As*LPMO10A and -B in chitin degradation, *Al. salmonicida* gene deletion strains were included in the cultivation experiments. The two single LPMO deletion strains showed a moderate decrease of the growth rate compared to the wild type, displaying a 30% increase in generation time (Fig. 8A and Table 1). In contrast to the biochemical assays that showed stronger synergy between recombinant *As*Chi18A and *As*LPMO10B compared to *As*LPMO10A, the cultivation assays showed that deletion of the single LPMOs resulted in the same growth reduction as deletion of both LPMOs. Deletion of the *AsChi18A* gene decreased growth to a larger extent than observed for the LPMO mutant strains (Fig. 8A), indicating that *As*Chi18A is more important than the LPMOs for the ability of *Al. salmonicida* to utilize chitin as a carbon source. The triple deletion mutant (ΔAΔBΔChi) was least able to utilize chitin as a source of nutrients, which also was clear from an agar-plate chitin solubilization assay where only a marginal disappearance of chitin was observed (Fig. S5). Growth of ΔAΔBΔChi and wild type on LB25 medium was on the other hand similar (Fig. S6), indicating that the gene deletions only influenced chitin utilization and not metabolism in general.

**Table 1.**
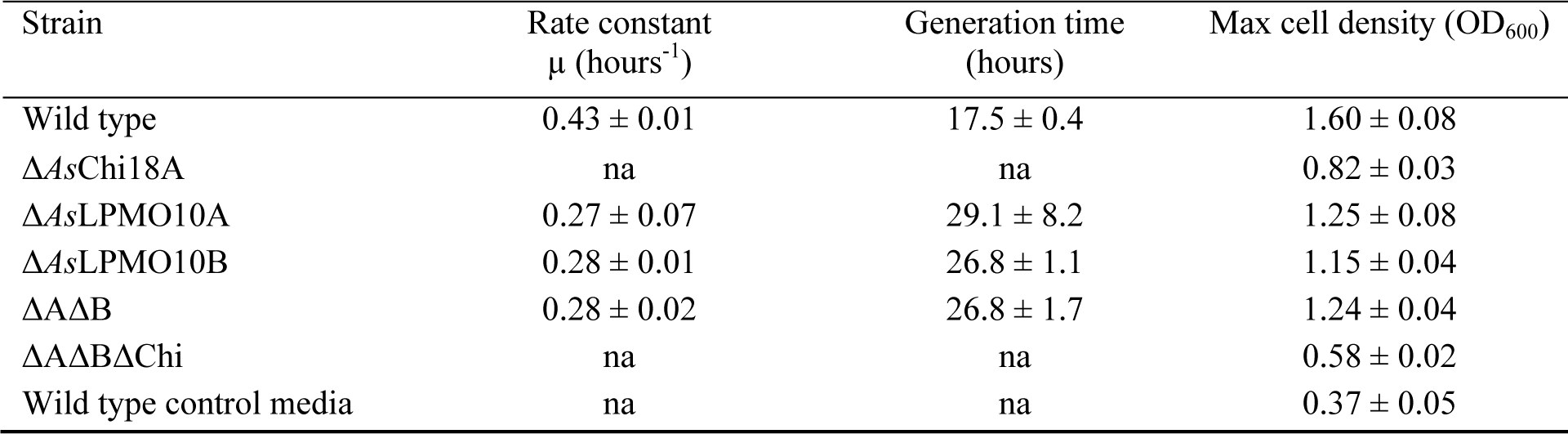
Growth rate and max cell density of *Al. salmonicida* and derivative mutant strains.

**Figure 8.**
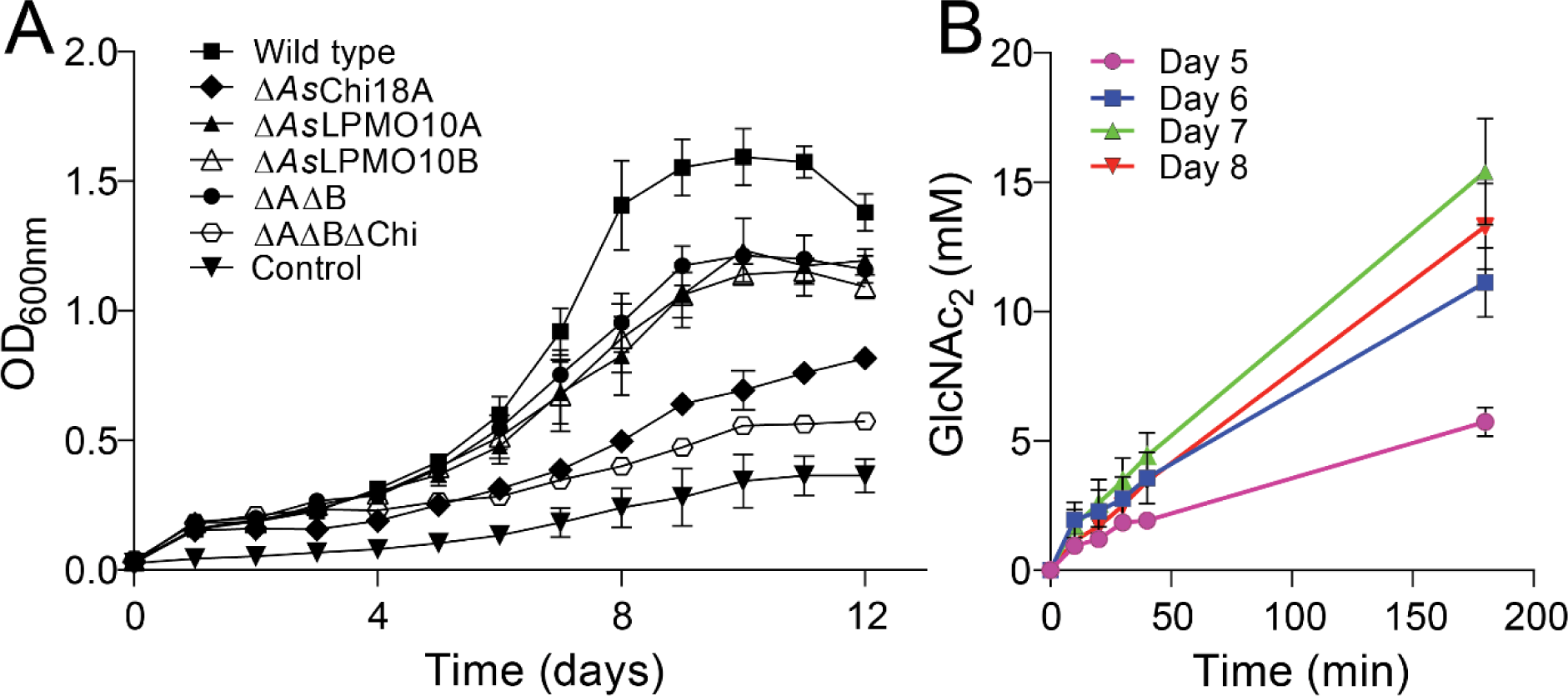
Growth of *Al. salmonicida* LFI1238 and derivate gene-deletion strains on β-chitin. (A) The growth of *Al. salmonicida* LFI1238 at 12 °C in minimal media supplemented with 1 % β-chitin. (B) Chitinase activity in the culture supernatant of *Al. salmonicida* growing on β-chitin. The chitinase activity was assayed by mixing a sample of the culture supernatant sampled at various time points with 15 mM chitopentaose and quantifying the (GlcNAc)_2_ resulting from hydrolysis over a period of 180 minutes. Error bars indicate standard deviation (n =3).

It should be noted that the wild type bacteria incubated in the minimal medium (Asmm) without added chitin obtained growth to OD 0.37±0.05 after 7 days incubation (Fig. 8 panel A and Table 1) due to the presence essential amino acids and traces of the LB25 pre-culture medium. Furthermore, it can also be observed that all bacterial strains incubated in the defined media supplemented with chitin increased ∼0.1 in OD within the first 24 hours. This is most likely caused by the presence of chitin monomers, dimers, oligosaccharides or other nutrients in the chitin substrate that could be utilized by the bacteria without the need of the chitinase or LPMOs.

To evaluate whether growth of the bacterium correlated with chitinolytic activity, the culture supernatant of wild type growing on β-chitin was sampled once a day in the period of highest growth (days 5-8) and analyzed for hydrolytic activity towards the soluble chitooligosaccharide, chitopentaose. Indeed, the chitin hydrolytic potential of the culture supernatant increased from day 5 to day 8 (Fig. 8B), indicating secretion of one or more chitinases (only dimeric and trimeric products were observed; large concentrations of GlcNAc would indicate the presence of a secreted *N*-acetylhexosaminidase).

### Gene expression analysis by PCR amplification of cDNA

Encouraged by the biochemically functional chitinolytic machinery of *Al. salmonicida* and the ability of the bacterium to metabolize chitin degradation products and chitin particles, it was of interest to couple these traits to transcription of genes representing the enzymes in the chitinolytic machinery. The pseudogene encoding parts of a family GH18 chitinase (*VSAL_I0902*; *As*Chi18B_p_) was also included in the analysis. RNA was isolated from *Al. salmonicida* LFI1238 grown on glucose, GlcNAc, (GlcNAc)_2_ and β-chitin (same cultures as shown in Fig. 1 and 8), from both exponential and stationary phase. Gene expression was assessed qualitatively by agarose gel chromatography (Table 2). The gene expression was assessed as positive if the target gene was amplified in two out of three biological replicates and at the same time no amplification was observed in PCR samples obtained in the control reactions having no reverse transcriptase during cDNA synthesis (examples shown in Fig. S7). The resulting data indicated that *AsChi18A*, *AsLPMO10B* and, surprisingly, the chitinase pseudogene, *AsChi18B_p_*, were expressed in the exponential phase during growth on all carbon sources. Similarly, expression of *AsChi18A* and *AsLPMO10A* were detected in the stationary phase, however not in all conditions. Expression of *AsLPMO10B* was only detected in the exponential phase during growth on GlcNAc.

**Table 2.** Gene expression of AsChi18A, AsLPMO10A, AsLPMO10B and AsChi18B_p_. Exp = Exponential phase, Stat.= Stationary phase. Data shown as positive (“+” on green background) or negative (“-“ on blue background) detection of expression, based on three biological replicates. *AsChi18A (VSAL_I0757), AsLPMO10A (VSAL_II0134), AsLPMO10B (VSAL_II0217)* and *AsChi18B****_p_*** *(VSAL_I0902*).

|  | GlcNAc |  | (GlcNAc)<sub>2</sub> |  | Glucose |  | β-Chitin |  |
| --- | --- | --- | --- | --- | --- | --- | --- | --- |
|  | Exp. | Stat. | Exp. | Stat. | Exp. | Stat. | Exp. | Stat. |
| *AsChi18A* | + | + | + | - | + | + | + | + |
| *AsLPMO10A* | + | + | + | + | + | - | + | - |
| *AsLPMO10B* | + | - | - | - | - | - | - | - |
| *AsChi18B<sub>p</sub>* | + | - | + | - | + | - | + | - |

### Proteomic analysis of expressed carbohydrate active enzymes (CAZymes)

To obtain a more complete understanding of chitin degradation by *Al. salmonicida* during growth, label free quantitative proteomics was used to identify and quantify proteins secreted by the bacterium when growing on this insoluble polysaccharide. Guided by the gene expression analysis (Table 2), cultures were grown to exponential phase on 1% β-chitin before harvesting and separation into supernatant and cell pellet fractions for analysis of both secreted and intracellular proteins. For analysis of bacteria and proteins binding to chitin, chitin from the growing culture was collected and boiled directly in sample buffer. These samples are referred to as “chitin-bound” samples and are enriched in proteins with high affinity for chitin. In total, 1179 proteins were identified (Supplementary data file 1), from which 20 were annotated as CAZymes, including glycoside hydrolases, transferase activities, lipid biosynthesis, glycogen metabolism, peptidoglycan (murein) and carbohydrate metabolic processes (Fig. 9, Table S2). In more detail, both LPMOs (*As*LPMO10A and *As*LPMO10B) and *As*Chi18A were identified, albeit not in all samples and at variable intensities. *As*LPMO10A was present at highest abundance amongst the CAZymes, especially in the chitin-bound samples. The protein was identified in all three biological replicates in all sampled conditions except in the bacterial pellet obtained from growth on glucose, where the protein only was identified in one biological replicate (Fig. 9).

**Figure 9.**
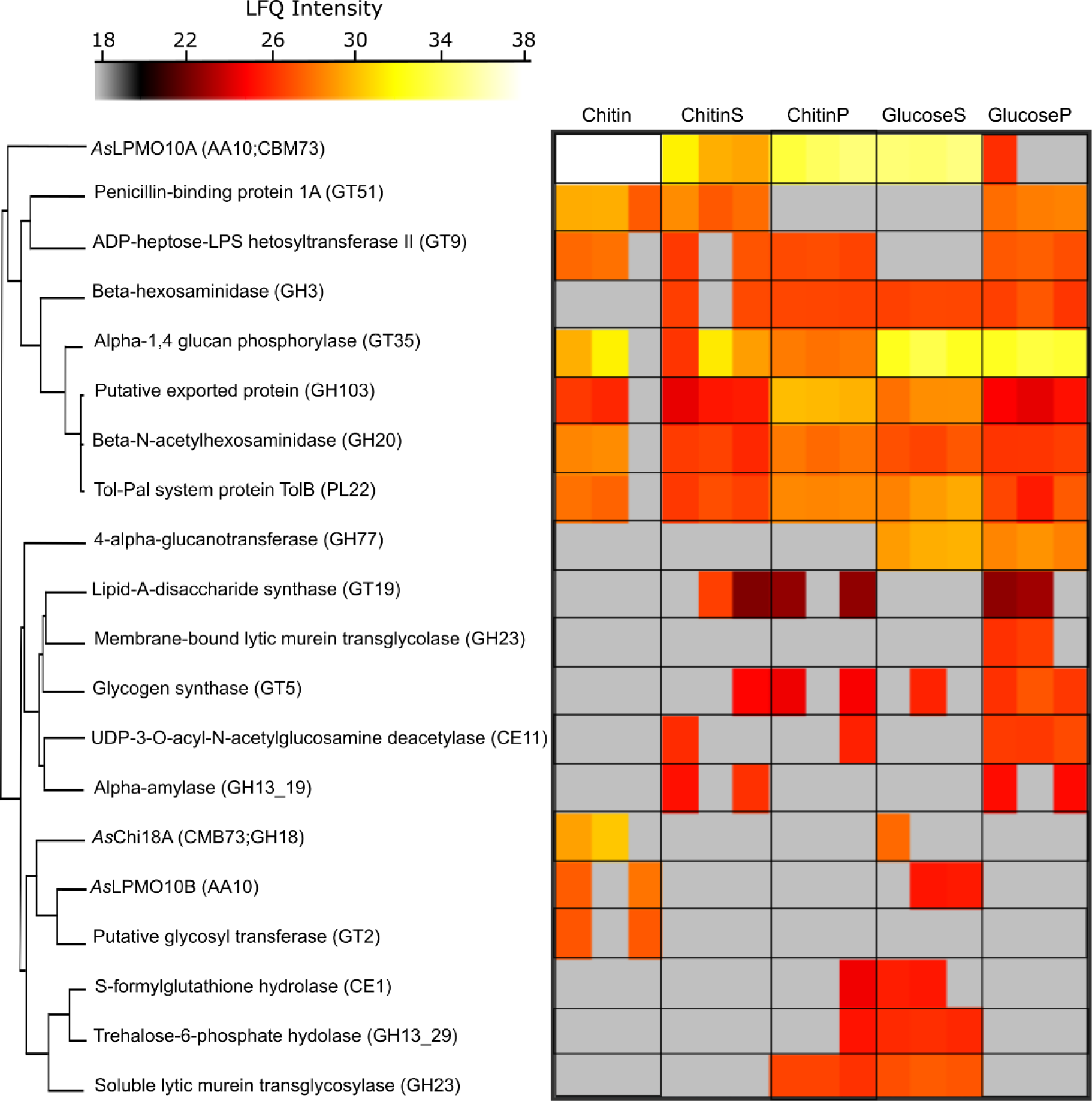
CAZymes expressed by *Al. salmonicida* LFI1238. Heatmap presentation of identified CAZymes and label free quantification intensities ranging from low intensity (grey), medium intensity (red) to high intensity (white). The data is presented as three biological replicates. Conditions are as following: proteins eluted from chitin obtained from the culturing experiment (Chitin), culture supernatant proteins from the chitin cultivation experiment (ChitinS), proteins extracted from the bacterial cells obtained from the chitin cultivation experiment (ChitinP), culture supernatant proteins obtained from culturing the bacterium on glucose (GlucoseS) and proteins extracted bacterial cell pellet from the glucose cultivation experiment (GlucoseP).

*As*Chi18A and *As*LPMO10B were only detected in the culture supernatant in one or two of the biological replicates obtained from growth on glucose, and in two out of three replicates of the chitin-bound samples. *As*Chi18A was only identified in the chitin-bound sample and the culture supernatant of the glucose grown samples. However, the chitinase was found at noticeable higher intensity in the chitin-bound samples compared to the supernatant samples obtained from cultivation on glucose.

Importantly, a GH20 β-*N*-acetylhexosaminidase (Uniprot ID: B6EGV7) was identified amongst the CAZymes. All samples showed a relatively similar abundance of this GH20. This enzyme, also called Chitobiase, is vital for hydrolyzing (GlcNAc)_2_ into two GlcNAc units, but also has the ability to depolymerize longer chitooligosaccharides (even aldonic acid chitooligosaccharides resulting from LPMO activity) (53). Sequence analysis revealed 58% identity between the *Al. salmonicida* GH20 identified (∼100% sequence coverage) and the biochemically characterized β-*N*-acetylhexosaminidase *Vh*NAG1 from *Vibrio harvey* 650 (54). The amino acids involved in catalysis and substrate binding are conserved (Fig. S8) indicating a function of the *Al. salmonicida* GH20 in chitin catabolism. It should be noted that *N,N*-diacetylchitobiose phosphorylases also can perform a role similar to β-*N*-acetylhexosaminidases. Interestingly, a family 3 glycosyl hydrolase (GH3), annotated as beta-hexosaminidase was also identified. GH3s have a broad range of substrate specificities, which mostly involves peptidoglycan recycling pathways. However, the marine bacteria *Pseudoalteromonas piscicida*, *Vibrio furnissi* and *Thermotoga maritima* encode GH3s that are believed to participate in intracellular chitin metabolism (55–57). The *As*GH3 enzyme was detected at similar levels in both glucose and chitin cultures, indicating that it is not dependent on chitin degradation. Also, the amino acid sequence of *As*GH3 was similar to the NagZ enzymes of this GH family (e.g. 67% sequence identity to NagZ of *V. cholerae*), which removes β-*N*-acetylglucosamine from ends of peptidoglycan fragments (58). 4-alpha-glucanotransferase (GH77) and membrane-bound lytic murein transglycosylase (GH23) were only detected when the bacterium was grown on glucose. A putative glycosyl transferase family 2 (GT2) was only detected in the chitin substrate fraction. GTs are generally involved in biosynthesis by transferring sugar moieties from activated donor molecules to specific acceptor molecules, forming glycosidic bonds.

### Analysis of the chitin catabolic pathway in *Al. salmonicida*

To assess the chitin catabolic pathway used by the bacterium, the proteomics data were scrutinized with the aim of identifying expressed proteins with a putative role in uptake, transport or downstream processing of chitin degradation products. An illustration of relevant findings and the suggested pathway is shown in Fig. 10. Guided by the biochemical assays and cultivation experiments, secreted *As*Chi18A, *As*LPMO10A and *As*LPMO10B are indicated to hydrolyze and cleave chitin into smaller oligosaccharides. It must be noted that *As*Chi18Bp, *As*Chi19Ap and *As*Chi18Cp are illustrated in context with *As*Chi18A based on conserved domains, rather than evidence of participating in extracellular hydrolysis of chitin. Interestingly, the chitinase pseudogene, *As*Chi18Bp, is one of few proteins exclusively identified in chitin samples. The GH20 β-*N*-acetylhexosaminidase, which shows a ∼3 fold increase in abundance during growth on chitin compared to glucose (p=0.0082, paired two-tailed t-test; Fig. S9), is indicated to hydrolyze (GlcNAc)_2_ into GlcNAc in the periplasmic space.

**Figure 10.**
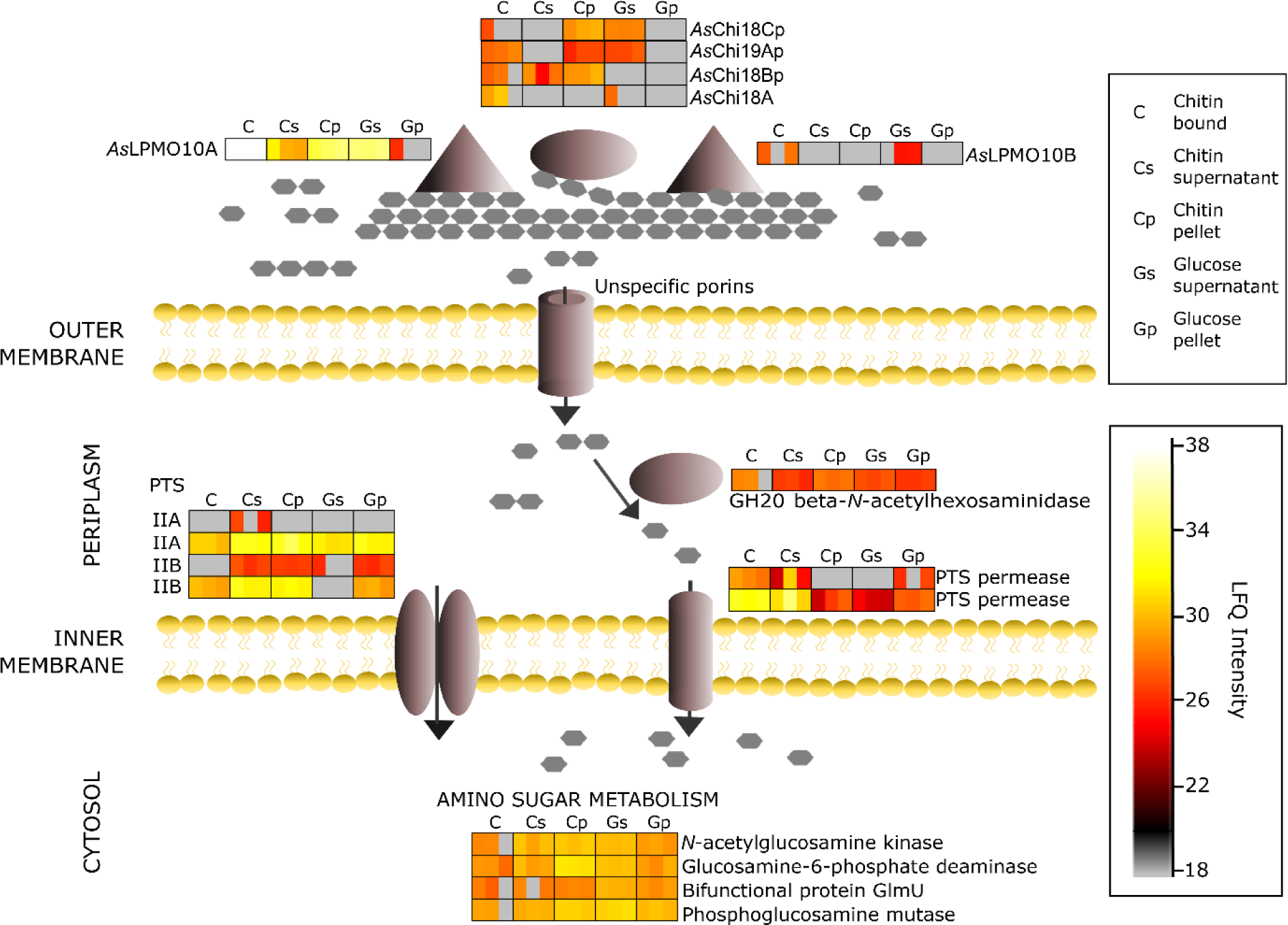
Putative chitin utilization pathway by *Al. salmonicida LFI238*. Illustration of detected proteins by label-free proteomics, aligned with their putative roles in the utilization pathway and the MaxLFQ Intensities. Enzymes acting on chitin: *As*Chi18A (B6EH15), *As*LPMO10A (B6EQB6), *As*LPMO10B (B6EQJ6), putative pseudogene chitinases (VSAL_I1414, VSAL_I1942, VSAL_I0902). Transport across membranes: Phosphotransferase system (PTS) component IIA (B6EGW5), PTS permease for *N*-acetylglucosamine and glucose (B6EHL6), PTS system, Lactose/Cellobiose specific IIB subunit (B6EMG0), PTS system permease for *N*-acetylglucosamine and glucose (B6ERZ1). Hydrolysis of (GlcNAc)_2_ into GlcNAc: beta-*N*-acetylhexosaminidase GH20 (B6EGV7), Amino sugar metabolism: Glucosamine-6-phosphate deaminase (B6EN78), UDP-*N*-acetylglucosamine pyrophosphorylase (B6EHG2), Phosphoglucosamine mutase (B6END8), *N*-acetyl-D-glucosamine kinase (B6EKQ4). A bar chart comparing the log2 LFQ values of the putative chitinolytic enzymes is shown in Fig. S9.

Utilization of extracellular sugars requires uptake and transportation across both the outer and inner membranes. With the lack of a functional chitoporin, other proteins relevant for outer membrane transport were investigated. Of proteins related to transport through the outer membrane, 14 proteins were identified, including outer membrane assembly factors and outer membrane proteins of the OmpA family, OmpU, TolC. These proteins are not generally known for sugar transport but cannot be excluded. For transport of sugars across the inner membrane, the most relevant transporters identified were 9 proteins assigned to the phosphoenolpyruvate-dependent sugar phosphotransferase system and two *N*-acetylglucosamine and glucose permeases (NagE). The latter transporters are likely contributing to translocation of GlcNAc across the inner membrane and showed increased abundance in chitin samples compared to glucose (Fig. 10). Two PTS component IIA and two Lactose/Cellobiose specific IIB subunits where identified, of which the lactose/cellobiose specific subunits likely contribute to sugar transportation across the inner membrane, were found upregulated during growth on chitin compared to glucose. Furthermore, out of 9 ABC transporter proteins identified, the four components not related to iron or amino acid transport were assessed. The ABC transport protein, “ATP binding component” (B6EMA3) shows increased abundance in the chitin-bound samples, whereas “ATP-binding protein” (B6ESL1) was only identified during growth on chitin. However, it is uncertain whether these proteins are involved in transport of GlcNAc/(GlcNAc)_2_. It should be noted that no ABC transporter proteins specific for (GlcNAc)_2_, or GlcNAc specific subunits could be identified, although these are common in transport of such sugars (59–61).

In terms of downstream processing of GlcNAc, the monosaccharide is most likely converted into GlcNAc6P by the permease NagE or *N*-acetylglucosamine kinase NagK (Fig. 10). N-acetylglucosamine deacetylase is encoded by the genome of *Al. salmonicida*, albeit was not identified in this experiment. Deacetylation of GlcNAc6P would result in GlcN-6P, a product further processed into Fru-6P by glucosamine-6-phosphate deaminase, an enzyme which was found at higher abundance in the chitin pellet samples compared to glucose (Fig. 10). Alternatively, GlcN-6P can be processed (in three steps) by Phosphoglucosamine mutase (EC 5.4.2.10), the bifunctional protein GlmU (*N*-acetylglucosamine-1-phosphate uridyltransferase (EC 2.3.1.157) and UDP-*N*-acetylglucosamine pyrophosphorylase (EC 2.7.7.23) into UDP-GlcNAc, a sugar that can be processed to other UDP sugars or utilized in pathways such as lipopolysaccharide biosynthesis or peptidoglycan synthesis. These enzymes were found in all conditions analyzed (Fig. 10).

## DISCUSSION

Knowing whether *Al. salmonicida* is able to utilize chitin as a source of carbon (and nitrogen) is important for understanding the ecology of the bacterium and its implications for pathogenicity. The literature contains conflicting information about this topic, but in the present study, we clearly demonstrate that *Al. salmonicida* is capable of degrading chitin to soluble chitooligosaccharides and to utilize these as a nutrient source. This capability is dependent on the single chitinase in the *Al. salmonicida* genome, despite the low *in vitro* activity of chitinase, and the ability of the LPMOs to degrade chitin. In the absence of *As*Chi18A, only products from LPMOs activity will be available to the bacterium. These products are oxidized chitooligosaccharides with a high degree of polymerization, that most likely cannot be taken up by the bacterium due to the absence of a specific outer membrane transporter (chitoporin). The fact that minor growth of the bacterium still is achieved in the absence of the chitinase is most likely due to the presence of a GH20 *N*-acetylhexosaminidase in the culture supernatant, that can depolymerize LPMO-generated chitooligosaccharides to GlcNAc, which can be taken up and catabolized by the bacterium. Another explanation may be that the chitooligosaccharides are cleaved by secreted pseudo-chitinases, proteins indeed observed by the proteomics data. In support for the latter hypothesis, minor growth on β-chitin and indications of degradation of colloidal chitin was observed for the *Al. salmonicida* ΔAΔBΔChi variant (Figs 8 and S5, respectively). Notably, the importance of a single chitinase for growth on chitin is not unique to *Al. salmonicida* LFI1238. In *C. japonicus, Cj*Chi18D is essential for the degradation of α-chitin despite the expression of three additional chitinases and two LPMOs (50). Similarly, a systematic genetic dissection of chitin degradation and uptake in *Vibrio cholerae* found the chitinase ChiA2 critical for growth on chitin, but not sufficient alone (62).

Both *As. salmonicida* LPMOs are required for obtaining maximum growth on chitin, an observation that is different than for the efficient chitin degrader *C. japonicus* where deletion of the chitin-active LPMO only resulted in delayed growth, but did not affect growth rate (50). This may be explained by the 50-fold lower activity of *As*Chi18A compared to *Cj*Chi18D of *C. japonicus*. In the latter organism, the contribution of the LPMOs in chitin solubilization is most likely minor compared to *Al. salmonicida*, for which the rate of depolymerization is almost equal for the LPMOs and the chitinase. *As*LPMO10A and -B are distinctly different in domain organization and sequence and the former enzyme is more active towards β-chitin than the latter. This may be related to the chitin binding properties of the enzymes as *As*LPMO10A binds better to both α- and β-chitin than *As*LPMO10B (Fig. 4). Alternatively, the difference in activity can be related to the ability of the components in the reaction mixture to generate reactive oxygen species such as hydrogen peroxide, e.g. by the oxidase activity of LPMOs as shown in several studies (63–65). In such a scenario, the discovery that LPMOs can use H_2_O_2_ as a co-substrate, and that the concentration of H_2_O_2_ in solution may be rate limiting for LPMO reactions (13, 66, 67), may account for activity differences between LPMOs when no external H_2_O_2_ is added to the enzyme reaction (only reductant).

The contribution of the LPMOs for chitin utilization by *Al. salmonicida* is most likely related to the synergy obtained when combining the LPMOs with the chitinase. Such synergy can be explained by the ability of *As*LPMO10s to cleave chitin chains that are inaccessible to *As*Chi18A (i.e. in the crystalline regions of the substrate). The newly formed chitin chain ends formed by LPMO activity, represent new points of attachment for the chitinases, thereby increasing substrate accessibility. Indeed, several studies have demonstrated this phenomenon (16, 68–70), including a study on the virulence-related LPMO from *Listeria monocytogenes* (71).

A surprising observed was made when combining both LPMOs and the chitinase in a chitin degradation reaction (Fig. 7, panels B&D). Here, no synergy was observed for β-chitin degradation and a lower than theoretical yield was obtained for α-chitin. This was unexpected since the bacterial cultivation assay indicated a cooperative relationship between the LPMOs as the reduced growth observed for two single LPMO deletion strains were similar to that observed for the double LPMO mutant strain (*As*ΔLPMO10A-ΔLPMO10B). The explanation for the lack of synergy is not straightforward, but it may be that a total concentration of 2 μM LPMO is too much for these reactions, giving rise to less bound enzyme to the substrate and thereby production of harmful reactive oxygen species (ROS) by the non-bound LPMO molecules. It is well established that LPMOs not bound to the substrate are more prone to autooxidation (13, 43, 72). Another explanation could be that a non-optimal enzyme stoichiometry could create competition for substrate binding sites. Indeed, Both LPMOs were expressed during growth on β-chitin, although *As*LPMO10A was detected in substantially higher abundance. As a matter of fact, *As*LPMO10A was the protein showing the highest abundance among the detected CAZymes, also when the bacterium was cultivated on glucose. This could imply that this LPMO has additional functions (this is discussed in more detail below). All three chitinolytic enzymes were observed in highest abundance in the samples obtained from the chitin particles, indicating high affinity of the enzymes towards chitin, a trait corroborated by the substrate binding experiments.

The proteomic analysis identified peptides from three pseudogenes. Interestingly, *As*Chi18Bp was only identified during growth on chitin, in contrast to the gene expression analysis where it was detected during growth in all carbon sources. This suggests a regulatory mechanism of translation influenced by the presence of chitin particles and that the relevant transcription factor regulating this gene still is functional. It is not uncommon that bacterial pseudogenes are expressed (73, 74) and Kuo & Ochman have hypothesized that this may be related to the regulatory region of the pseudogenes still remaining intact (74). It must be noted that translation of a pseudogene does not necessarily equal a functional protein. Indeed, our data showing a large growth impairment upon *AsChi18A* deletion suggest that translation of pseudogenes is insufficient for chitin degradation, although, as previously noted, a minor growth also can be observed for the triple knock out strain. Pseudogenes have long been considered to only represent dysfunctional outcomes of genome evolution, and the multitude of pseudogenes in *Al. salmonicida* LFI1238 possibly reflects its adaption to a pathogenic lifestyle. On the other hand, there is increasing evidence indicating that pseudogenes can have functional biological roles, and recent studies have shown that pseudogenes potentially regulate expression of protein-coding genes (reviewed in (75, 76)).

An intriguing observation of chitin catabolism by *Al. salmonicida* is the absence of key regulatory proteins such as ChiS and Tfox in the proteomics data. These regulatory proteins are important for chitin catabolism in other bacterial species in the *Vibrionacea* (18, 31, 33, 34). There is no doubt that *Al. salmonicida* is capable of chitin catabolism, thus the bacterium may have evolved an alternative mechanism for regulating the chitin utilization loci. In support of this hypothesis, the gene encoding the periplasmic chitin binding protein, which activates ChiS when bound to (GlcNAc)_2_ (31), is disrupted in the *Al. salmonicida* genome (29).

Although the *Al. salmonicida* chitinolytic system clearly is active and functional, there are some observations that may indicate other or additional functions of the chitinolytic enzymes. Firstly, the activity of the chitinase is substantially lower that what would be expected for an enzyme dedicated to chitin hydrolysis. Secondly, the dominantly expressed LPMO (*As*LPMO10A) is not essential for chitin degradation and is also abundantly expressed when the bacterium is cultivated on glucose. These observations could be associated with the adaption of a pathogenic lifestyle where the need for chitin as a nutrient source has been reduced, but could also indicate other or additional functions, as for example roles in virulence. The notion of chitinases having additional functions has been shown in several studies, for example cleavage of mucin glycans by the *V. cholerae* chitinase Chi2A (77) and hydrolysis of LacdiNAc (GalNAcβ1-4GlcNAc) and LacNAc (Galβ1-4GlcNAc) by the *L. monocytogenes* and *Salmonella typhimurium* chitinases (78). Such substrates were not evaluated by activity assays with *As*Chi18A. Moreover, incubation of *As*Chi18A with mucus collected from Atlantic salmon skin revealed an unidentifiable product (different from the negative control), but determination of its identity was unsuccessful.

Compared to other virulence related chitinases, *As*Chi18A has a similar size, but different modular architechture. For example, ChiA2 from *V. cholerae*, which has been shown to improve survival of the bacterium in the host intestine, also contains around 800 amino acids, but the GH18 domain is close to the N-terminus and a CBM44 and a CBM5 chitin-binding domain are present on the C-terminal side. As already noted, ChiA2 has been shown to cleave intestinal mucin (releasing GlcNAc), but has a deep substrate binding cleft and resembles an *exo*-chitinase (85% sequence identity to the structurally resolved *exo*-chitinase of *Vibrio harveyi*; (79)). An unusual property of *As*Chi18A is its double pH optimum, shown by enzyme activity being approximately equal at pH 4 and 7 (Fig. 5B). Chitinases usually display a single pH optimum, but double pH optima are not uncommon for hydrolytic enzymes, e.g. like phytase from *Aspergillus niger* (80) and β-galactosidase from *Lactobacillus acidophilus* (81). It is possible that this property is associated with the chitinase being utilized in environments that vary in pH. If the *Al. salmonicida* chitinase has evolved an additional role than chitin degradation, the same question applies for the LPMOs. Both LPMOs are active towards chitin, but it is not certain that this is the intended substrate of these enzymes. For instance, GbpA, an LPMO from *V. cholera*, has activity towards chitin (53), but its main function seems to be related to bacterial colonization of transfer vectors (e.g. zoo-plankton), the host epithelium (e.g. human intestine) or both (82, 83). The LPMO of *L. monocytogenes* is also active towards chitin (71), but the gene encoding this enzyme is not expressed when the bacterium grows on chitin (on the other hand, the *L. monocytogenes* chitinase-encoding genes are expressed when the bacterium is grown on chitin (71, 84)). The LPMO of the human opportunistic pathogen *Pseudomonas aeruginosa*, CbpD, was recently shown to be a chitin-active virulence factor that attenuates the terminal complement cascade of the host (85). In the present study, both LPMOs were expressed in the presence of chitin, but also in the glucose control condition, indicating that regulation is not controlled by chitin or soluble chitooligosaccharides. Thus, chitin may represent a potential substrate for these LPMOs, but possibly not the (only) biologically relevant substrate.

On the other hand, some LPMOs are designed to only disrupt and disentangle chitin fibers, rather than to contribute to their degradation in a metabolic context, namely the viral family AA10 LPMOs (also called spindolins) (41). These LPMOs are harbored by insect-targeting entomopox- and baculoviruses, and have been shown to disrupt the chitin containing peritrophic matrix that lines the midgut of insect larvae (86). The main function proposed for the viral LPMOs is to destroy the midgut lining in order to allow the virus particles to access the epithelial cells that are located underneath. Since the scales and gut of fish are indicated to contain chitin (5, 6), it is tempting to speculate that the role of the fish pathogen LPMOs is similar to that of viral LPMOs, namely to disrupt this putatively protective chitin layer in order to provide an entry point to the bacteria for infection.

In conclusion, the present study shows that *Al. salmonicida* LFI1238 can degrade and catabolize chitin as a sole carbon source, despite possessing a chitinolytic pathway assumed to be incomplete. Our findings imply that the bacterium can utilize chitin to proliferate in the marine environment, although possibly not as efficient as other characterized chitinolytic marine bacteria. Nevertheless, it is likely that this ability can be of relevance for the spread of this pathogen in the ocean. Finally, our discovery that pseudogenes are actively transcribed and translated indicates that such genes cannot be disregarded as being functionally important.

## METHODS AND MATERIALS

### Bacterial strains and culturing conditions

*Al. salmonicida* strain LFI1238 originally isolated from the head kidney of diseased farmed cod (*Gadhus morhua;* (*29*)) and mutant strains (see below) were routinely cultivated at 12 °C in liquid Luria Broth (LB) supplemented with 2.5% sodium chloride (LB25; 10 g/L tryptone, 5 g/L yeast extract, 12.5 g/L NaCl) or solid LB25 supplemented with 15 g/L agar powder (LA25), and if applicable 2% (w/v) colloidal chitin made from α-chitin (gift from Silje Lorentzen). Growth analysis was performed at 12 °C in *Al. salmonicida* specific minimal media (Asmm: 100 mM KH_2_PO_4_, 15 mM NH_4_(SO_4_)_2_, 3.9 μM FeSO_4_×7H_2_O, 2.5 % NaCl, 0.81 mM MgSO_4_×7H_2_O, 2 mM valine, 0.5 mM isoleucine, 0.5 mM cysteine, 0.5 mM methionine and 40 mM glutamate). Prior to inoculation of Asmm media, strains were grown up to 48 hours in 10-15 mL LB25 at 200 rpm. 1 mL bacteria were harvested by centrifugation at 6000 × *g* for 1 minute, followed by immediate resuspension of the pellet in 1 mL Asmm. The cell suspension was transferred to the final cultures by a 1:50 dilution in media supplemented with 0.2% glucose, 0.2% N-acetyl-D-glucosamine, 0.2 % diacetyl-chitobiose (Megazyme, Bray, County Wicklow, USA) or 1 % β-chitin from squid pen purchased from France Chitine (Batch 20140101, Orange, France). Culture volumes ranged from 5-50 mL. Final cultures were incubated at 12 °C with shaking at 175 rpm. Growth was measured by optical density (OD_600_) using Ultrospec® 10 Cell Density Meter (Biochrom). The baseline was set by using sterile Asmm media with or without 1% β-chitin. OD_600_ measurements of the β-chitin cultures was performed by allowing the cultures to settle for 30 seconds before collecting 1 mL for measurement.

### Generation of gene deletion strains

LFI1238 derivative in-frame deletion mutants *ΔAsChi18A, ΔAsLPMO10A, ΔAsLPMO10B, ΔAsLPMO10A-ΔLPMO10B and ΔLPMO10A-ΔLPMO10B-ΔChi18A* (also referred to as ΔAΔBΔChi) were constructed by allelic exchange as described by others (87, 88). For clarification, Table 3 lists the target genes, their associated protein name, predicted carbohydrate-active enzyme family (CAZyme family) and corresponding CAZyme annotated name applied throughout this study.

**Table 3.**
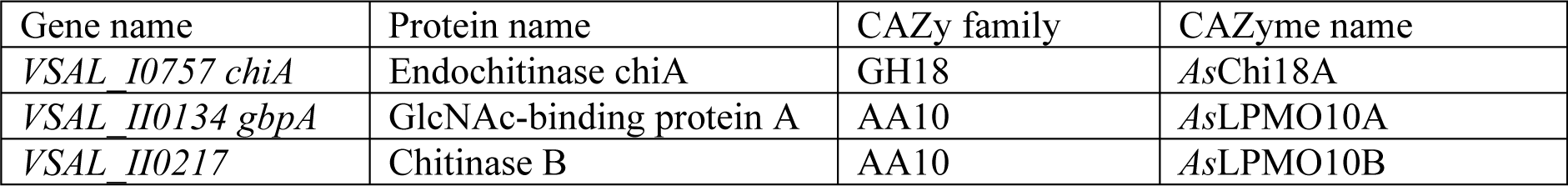
Description of genes targeted for deletion.

| Gene name | Protein name | CAZy family | CAZyme name |
| --- | --- | --- | --- |
| *VSAL_I0757 chiA* | Endochitinase chiA | GH18 | AsChi18A |
| *VSAL_II0134 gbpA* | GlcNAc-binding protein A | AA10 | AsLPMO10A |
| *VSAL_II0217* | Chitinase B | AA10 | AsLPMO10B |

Primers were ordered from eurofins Genomics (Ebersberg, Germany), and designed with restriction sites and regions complementary to the pDM4 cloning vector to allow for in-fusion cloning. Table 4 lists primers used for construction of the deletion alleles. For construction of ΔAsChi18A, the flanking regions upstream and downstream of the *AsChi18A* gene were amplified using primer pairs GH18_YF/GH18_IR and GH18_IF/GH18_YR, respectively. The two PCR fragments were fused by overlapping extension PCR where complementarity in the 5′ regions of the primers resulted in linkage of the *AsChi18A* -flanking regions. Δ*AsLPMO10A* and Δ*AsLPMO10B* was constructed in the same manner as described for *ΔAsChi18A* using the listed primers (Table 4).

**Table 4.**
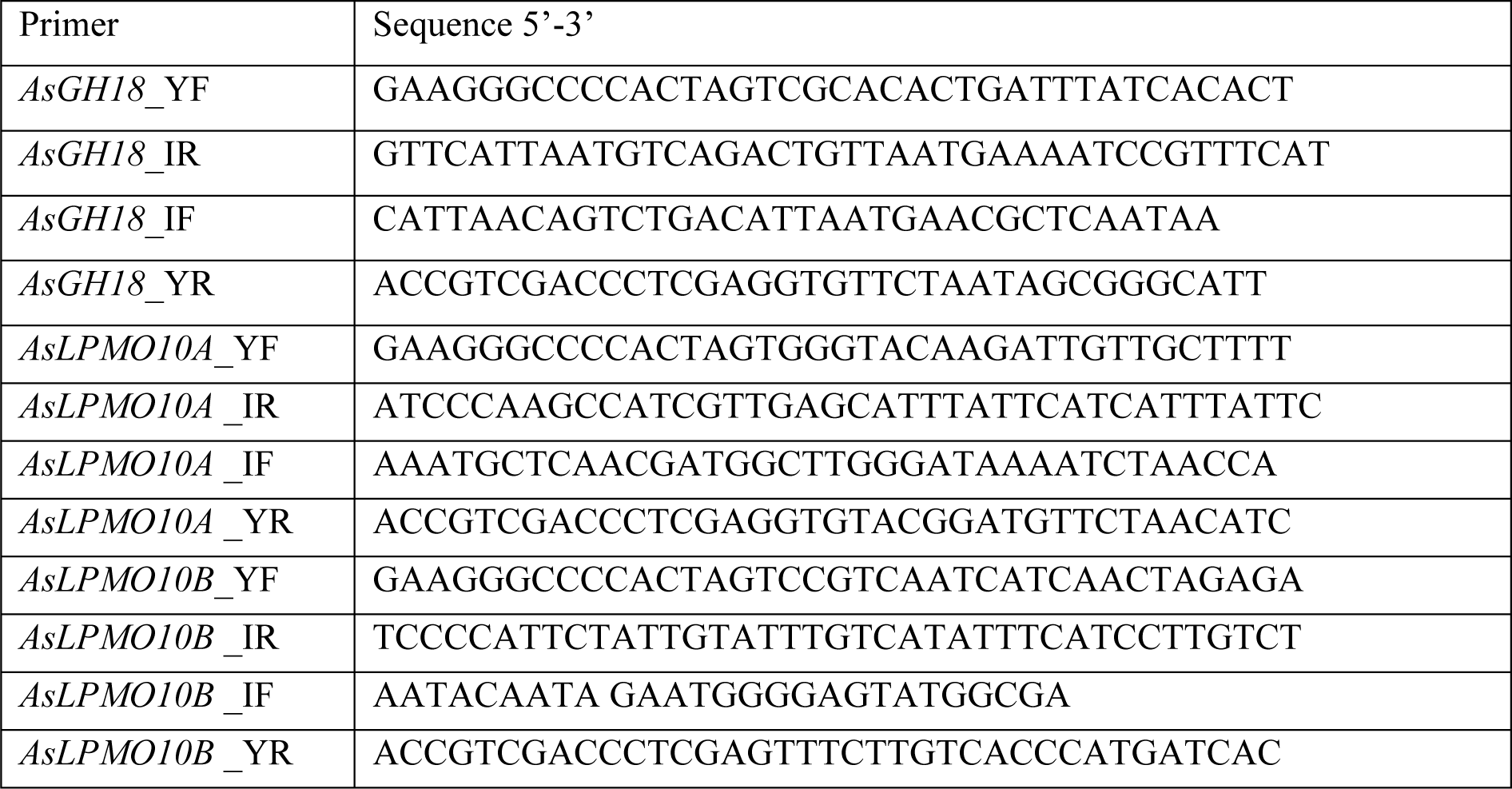
Primers used for construction of in frame deletion mutants.

The final PCR products were inserted into the suicide vector pDM4 by In-Fusion ® HD cloning (Takara Bio USA, Inc). In short, pDM4 linearized with SpeI and XhoI was gently mixed with 5× In-Fusion HD Enzyme premix, purified PCR fragment (purified using Nucleospin® Gel and PCR Clean-up, MACHEREY-NAGEL GmbH & Co. KG), and H_2_O to acquired final volume. Ratio of insert and linearized vector was determined using the online tool “In-Fusion molar ratio calculator” (Takara Bio USA, Inc). The reaction mix was incubated at 50 °C for 15 min. Following incubation, the reaction mix was placed on ice for 20 min and transformed into *E. coli* S17-1 λpir by standard transformation techniques.

Conjugation was performed as described by others (87–90). In brief, pelleted cells from 1 mL *E.coli* S17-1 donor cells (OD_600_ 0.60-0.80) and 1 mL *Al. salmonicida* LFI1238 recipient cells (OD_600_ 1.00-1.40) were washed in LB, mixed and transferred to LA1 as a spot. The spot plate was incubated 6 hours in room temperature and ∼17 hours at 12 °C. The next day, the cell spot was collected and resuspended in 2 mL LB25, grown for 24 hours with shaking and spread onto LA25 containing chloramphenicol (2 μL/mL). Potential transconjugates were re-streaked on LA25 2CAM, incubated for 3-5 days and tested for integration of the pDM4 construct by colony PCR using a combination of primers annealing within and outside the integrated plasmid (Table S3). Next, confirmed transconjugates were grown in LB25 to OD_600_ 0.4 and spread onto LA25 containing 5% sucrose. Colonies appearing within 5 days were tested for excision of the integrated plasmid by sequentially patching single colonies onto LA25 plates containing 2CAM or 5% sucrose. Mutants showing loss of resistance to CAM and presence of gene-deletion product (colony PCR using primer pairs *AsΔChi18A*_For/ *AsΔChi18A*_Rev), was confirmed by GATC Biotech Sanger sequencing (Eurofins genomics, Germany).

Mutant strains containing multiple gene deletions were generated in a step-wise manner. Specifically, LFI1238ΔAsLPMO10A were recipient cells for pDM4-ΔAsLPMO10B. Similarly, the resulting ΔAsLPMO10A/ΔLPMO10B strain were recipient cells for pDM4-ΔAsChi18A, thus generating the triple mutant strain ΔLPMO10A/ΔLPMO10B/ΔChi18A. All strains and vectors are listed in Table 5.

**Table 5.** Complete list of strains and vectors.

| Strain or plasmid | Comment | Ref. |
| --- | --- | --- |
| LFI1238 | *Aliivibrio salmonicida* strain LFI1238 | § |
| S17-1 λpir | *Escherichia coli* conjugation donor strain S17-1 λpir | (102) |
| AsΔChi18A | LFI1238 containing gene deletion ΔChi18A | * |
| AsΔLPMO10A | LFI1238 containing gene deletion ΔLPMO10A | * |
| AsΔLPMO10B | LFI1238 containing gene deletion ΔLPMO10B | * |
| AsΔLPMO10A-Δ10B | LFI1238 containing gene deletions ΔLPMO10A and ΔLPMO10B | * |
| AsΔLPMO10A-Δ10B-ΔChi | LFI1238 containing gene deletions ΔLPMO10A, ΔLPMO10B and ΔChi18A | * |
| pDM4 | pDM4 SacB suicide plasmid/ cloning vector | (90) |
| pDM4-AsΔChi18A | pDM4-construct designed for allelic exchange and deletion of AsChi18A | * |
| pDM4-AsΔLPMO10A | pDM4-construct designed for allelic exchange and deletion of AsLPMO10A | * |
| pDM4-AsΔLPMO10B | pDM4-construct designed for allelic exchange and deletion of AsLPMO10B | * |

§Originally isolated by the Norwegian Institute of Fisheries and Aquaculture Research, N-9291 Tromsø,
Norway, but provided by Simen Foyen Nørstebø for this study. \*This study.

### Cloning, expression and purification

Codon-optimized genes encoding the following *As*LPMO10A (residues 1-491, UniProt ID; B6EQB6), *As*LPMO10B (residues 1-395, UniProt ID; B6EQJ6) and *As*Chi18A (residues 1-846, UniProt ID; B6EH15) from *Al. salmonicida* (LFI1238) were purchased from GenScript (Piscataway, NJ, USA). Gene-specific primers (Table 6), with sequence overhangs corresponding to the pre-linearized pNIC-CH expression vector (AddGene, Cambridge, Massachusetts, USA) were used to amplify the genes in order to insert them into the vector by a ligation independent cloning method (91). All the cloned genes contained their native signal peptides. Sequence-verified plasmids were transformed into ArcticExpress (DE3) competent cells (Agilent Technologies, California, USA) for protein expression. Cells harboring the plasmids were inoculated and grown in Terrific Broth (TB) medium supplemented with 50 µg/mL of kanamycin (50 mg/mL stock). Cells producing the full-length *As*LPMO10s were cultivated in flask-media at 37 °C until OD = 0.700, cooled down for 30 min at 4 °C, induced with 0.5 mM IPTG and incubated for 44 hours at 10 °C with shaking at 200 rpm. Cells producing *As*Chi18A were grown in a Harbinger LEX bioreactor system (Epiphyte Three Inc, Toronto, Canada) using the same procedure described above, although the cell were cultured for a shorter time period (12 hours) and air was pumped into the culture by spargers. Successively, cells were harvested using centrifugation and the periplasmic extracts were generated by osmotic shock (92). The periplasmic fractions, containing the mature proteins (signal peptide-free), were sterilized by filtration (0.2 µm) before purification (see below).

**Table 6.**
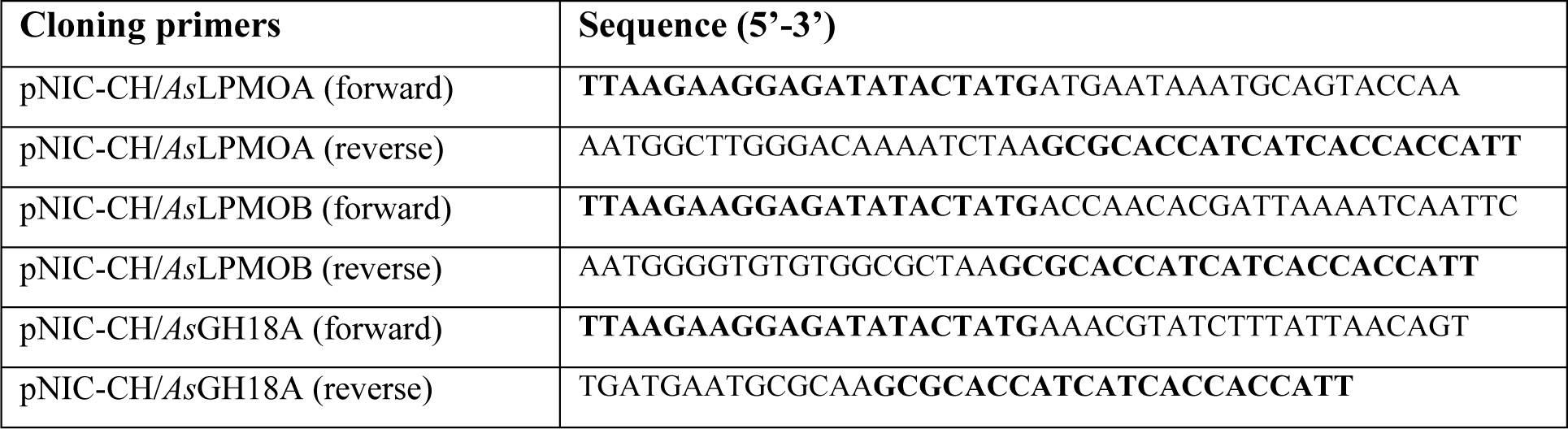
Cloning primers for *As*LPMO10A and -B and *As*Chi18A.

*As*LMO10A and *As*LPMO10B were purified by anion exchange chromatography using a 5 mL HiTrap DEAE FF column (GE Healthcare) followed by hydrophobic interaction chromatography (HIC) using a 5 mL HiTrap Phenyl FF (HS) column (GE Healthcare). For the ion exchange procedure, proteins in the periplasmic extract were applied on the column using a binding buffer containing 50 mM Bis-Tris-HCl pH 6.0. After all non-bound proteins had passed through the column, bound proteins were eluted by applying a linear gradient (0 to 100 % in 20 column volumes with a flow rate of 1 mL/min), using an elution buffer containing Bis-Tris-HCl pH 6.0 and 500 mM NaCl. Fractions were collected and analyzed for the presence of LPMO using SDS-PAGE. Fractions containing LPMO were pooled and adjusted to 1M (NH_4_)_2_SO_4_ and applied on the HIC column using a binding buffer consisting of 50 mM Tris-HCl pH 7.5 and 1 M (NH_4_)_2_SO_4._ Following elution of unbound proteins, bound proteins were eluted by applying a linear gradient (0 to 100% over 20 column volumes with a flow rate of 1.5 mL/min), using an elution buffer containing 50 mM Tris-HCl pH 7.5. In addition, *As*LPMO10B was further purified by size exclusion chromatography using a HiLoad 16/60 Superdex 75 column operated at 1 mL/min and with a running buffer containing 1X PBS, pH 7.4.

*As*Chi18A was purified by immobilized metal affinity chromatography using a HisTrap FF 5 mL column (GE Healthcare). The periplasmic extract containing *As*Chi18A was applied on the column using a binding buffer consisting of 20 mM Tris-HCl pH 8.0 and 5 mM imidazole, using a flow rate of 3 mL/min. Bound proteins were eluted from the column by applying a linear gradient (0 to 100 % over 20 column volumes with a flow rate of 3 mL/min) with an elution buffer containing 20 mM Tris-HCl pH 8.0 and 500 mM imidazole. Fractions containing the pure protein, identified by SDS-PAGE, were pooled and concentrated using Amicon Ultra centrifugal filters (Millipore, Cork, Ireland).

Protein purity was analyzed by SDS-PAGE. Concentrations of the pure proteins were determined by measuring A_280_ and using the theoretical molar extinction coefficients of the respective enzyme (calculated using the ExPASy ProtParam tool) to estimate the concentration in mg/mL. Before use, *As*LPMO10A and *As*LPMO10B were saturated with Cu(II) by incubation with excess of CuSO_4_ in a molar ratio of 1:3 for 30 minutes at room temperature. The excess Cu(II) was eliminated by passing the protein through a PD MidiTrap G-25 desalting column (GE Healthcare) equilibrated with 50 mM Tris-HCl pH 8.0 and 150 mM NaCl.

### Preparation of substrates

The substrates used in the assays were either squid pen β-chitin (France Chitin, Orange, France), shrimp shell α-chitin purchased from Chitinor As (Avaldsnes, Norway) and skin mucus of *Salmo salar*. Skin mucus was collected from freshly killed farmed Atlantic salmon purchased from the Solbergstrand Marine Research Facility (Drøbak, Norway). The mucus was gently scraped off the skin of the fish using a spatula and stored in plastic sample tubes at -20°C until use.

### Enzyme activity assays

For activity assays, chitin was suspended in 20 mM Tris-HCl pH 7.5, in 2 mL Eppendorf tubes to yield a final concentration of 10 mg/mL. All reactions were incubated at 30 °C and stirred in an Eppendorf Comfort Thermomixer at 700 rpm. For LPMO reactions, the final enzyme concentrations were 1 µM and reactions were started by the addition of 1 mM of ascorbic acid (this activates the LPMOs). Similar reaction conditions were used for *As*Chi18A, although the final enzyme concentration used was 0.5 µM and ascorbic acid was not added in the reactions. At regular intervals, samples were taken from the reactions and the soluble fractions were separated from the insoluble substrate particles using a 96-well filter plate (Millipore) operated with a vacuum manifold. Subsequently, the soluble fraction of *As*LPMO10s-catalyzed reactions were incubated with 1.5 µM of a chitobiase from *S. marcescens* (also known as *Sm*CHB or *Sm*GH20A) at 37 °C overnight in order to convert LPMO products (oxidized chitooligosaccharides of various degree of polymerization) to *N*-acetylglucosamine (GlcNAc) and chitobionic acid (GlcNAcGlcNAc1A) as previously described in (53, 93), followed by a sample dilution with 50 mM H_2_SO_4_ in a ratio of 1:1 prior quantification by HPLC (see below). The soluble fractions of *As*Chi18A reactions, were diluted with H_2_SO_4_ after the filtration step, which stopped the enzymatic reaction, before quantification of (GlcNAc)_2_ by HPLC (see below). Additionally, in order to collect samples for product profiling by matrix-assisted laser desorption/ionization time of flight mass spectrometry (MALDI-TOF MS, see below) of the two *As*LPMO10s-catalyzed reactions, 5 µL of the soluble fraction was sampled after filtration and kept at -20 °C prior to analysis.

### Analysis and quantification of native and oxidized chitooligosaccharides, (GlcNAc)_2_ and GlcNAc

Qualitative analysis of the native and oxidized products of the *As*LPMO10A and -B soluble fractions were performed by MALDI-TOF MS using a method developed by G. Vaaje-Kolstad et al. (12). For this analysis, 1 µL of sample was mixed with 2 µL 2,5-dihydroxybenzoic acid (9 g.L^-1^, prepared in 150:350 H_2_O/Acetonitrile), applied to a MTP 384 target plate in ground steel TF (Bruker Daltonics) and dried under a stream of warm air. The samples were analyzed with an Ultraflex MALDI-TOF/TOF instrument (Bruker Daltonics GmbH, Bremen, Germany) equipped with a Nitrogen 337 nm laser beam, using Bruker FlexAnalysis software. Quantitative analysis of all soluble products formed by the chitinolytic enzymes or GlcNAc or (GlcNAc)_2_ in culture supernatants was performed by ion exclusion chromatography using a Dionex Ultimate 3000 UHPLC system (Dionex Corp., Sunnyvale, CA, USA), equipped with a Rezex RFQ-Fast acid H^+^ (8%) 7.8% x 100 mm column (Phenomenex, Torrance, CA). The column was pre-heated to 85 °C and was operated by running 5 mM H_2_SO_4_ as a mobile phase at a flow rate of 1 mL/min. The products were separated isocratically and detected by UV absorption at 194 nm. The amount of GlcNAc and (GlcNAc)_2_ were quantified using standard curves. Pure GlcNAc and (GlcNAc)_2_ were obtained from Sigma and Megazyme, respectively. In order to quantify chitobionic acid (GlcNAcGlcNAc1A), a standard was produced in-house by treating chitobiose (Megazyme, Bray, Irleand) with a chitooligosaccharide oxidase (ChitO) from *Fusarium graminearum*, which yields 100% conversion of chitobiose to chitobionic acid, a method previously described by J. S. M. Loose et al. (53). Standards were regularly analysed in each run.

### Analysis of chitinase activity in culture supernatants

To analyze presence of chitinolytic activity in the supernatant of *Al. salmonicida* when growing on β-chitin, 1 mL sample of wild type bacterial culture was harvested at time points during growth on chitin. The sample was centrifugated and the supernatant filter sterilized using 0.22µm sterile Ultra-free centrifugal filters. 500 µL filter sterilized supernatant was concentrated to 100 µL using Amicon ultra centrifugal filter units with 3 000 Da cut-off (Merck Millipore, Cork Ireland) and washed three times in 10 mM Tris pH 7.5, 0.2 M NaCl (Tris-HCl NaCl). The concentrated supernatant containing secreted enzymes were stored in Tris-HCl at 4 °C until use. The presence of chitinolytic activity was assessed by mixing 100 µM chitopentaose with 15 µL enzyme cocktail in 20 mM Tris pH 7.5 0.2 M NaCl and incubated at 30 °C. The generated products were analyzed and quantified by ion exclusion chromatography as described above.

### Protein binding assays

The binding capacity of *As*LPMO10s and *AsChi*18A on α-chitin and β-chitin was tested, suspending 10 mg/mL of substrate in 20 mM Tris-HCl pH 7.5 to a total volume of 350 μL in 2 mL Eppendorf tubes. Reactions were started by the addition of *As*LPMO10A or –B (0.75 µM final concentration) or *AsChi*18A (0.50 µM), which were incubated in 2 mL Eppendorf tubes, at 30 °C and stirred in an Eppendorf Comfort Thermomixer at 700 rpm. Samples were taken (100 µL) after 2 hours and immediately filtrated using a 96-well filter plate (Millipore) operated with a vacuum manifold to obtain the unbound protein fraction. In order to assess the percentage of bound proteins to the substrate, control samples with only enzyme and buffer were performed, representing the maximum quantity of protein present in the samples (100%). The protein concentration in each sample was determined using the Bradford assays (Bio-Rad, Munich, Germany).

### RNA isolation and gene expression analysis

To analyze expression of specific genes as previously done by e.g. T. M. Wagner et al. (94), samples were taken at mid exponential phase (OD = 0.6-0.7) and early stationary phase (OD = 1.0-1.3).1 mL sample of each culture was directly transferred to 2 mL RNAprotect cell reagent (Qiagen, Hilden, Germany). The samples were vortexed 5 sec, incubated 5 min at room temperature and subsequently harvested by centrifugation at 4000 × *g*, for 10 min at 4 °C. The supernatant was carefully decanted, and the cell pellet stored at -20 °C until cell lysis and RNA isolation. RNA isolation was performed using Qiagen RNeasy Mini Kit (Qiagen, Hilden, Germany) using the Quick-Start protocol. In order to disrupt the bacterial cell wall before isolation, the pellet was lysed using 200 µL Tris-EDTA pH 8.0 supplemented with 1 mg/mL lysozyme, vortexed for 10 sec and subsequently incubated at room temperature for 45 min. 700 µL buffer RLT (kit buffer, Qiagen) supplemented with 10 µL/mL β-mercaptoethanol was added to the sample and mixed vigorously before proceeding with the protocol. The quantity of isolated RNA was determined using NanoDrop.

Residual genomic DNA (gDNA) was removed using The Heat&Run gDNA removal kit (ArcticZymes®, Tromsø, Norway). 8 µL of the RNA samples was transferred to a RNase free Eppendorf tube on ice. For each 10 µL reaction, 1 µL of 10× reaction buffer and 1 µL Heat-labile-dsDNase was added. The suspension was gently mixed and incubated at 37 °C for 10 min. To inactive the enzyme, samples were immediately transferred to 58 °C for 5 min. The RNA concentration was measured using the nanodrop before proceeding to copy DNA (cDNA) synthesis.

cDNA synthesis was performed using iScript™ Reverse Transcription Supermix (Bio-Rad, Hercules, CA, USA). For each sample, 100 ng RNA, 4 µL 5× iScript™ Reverse transcription Supermix and RNase free water to a total volume of 20 µL was assembled in PCR reaction tubes. All samples were additionally prepared with iScript™ no reverse transcriptase control supermix to account for residual gDNA in downstream analysis. The cDNA synthesis of the samples were performed by using a SimpliAmp™ Thermal Cycler (Thermo Fischer Scientific, USA) with the following steps: priming at 25 °C for 5 min, reverse transcription at 46 °C for 20 min, and inactivation of the reverse transcriptase at 95 °C for 1 min. The synthesized cDNA was stored at -20 °C until analysis.

The cDNA samples were screened for presence of *AsChi18A, AsLPMO10A, AsLPMO10B* and *VSAL_I0902*/*AsChi18Bp* by PCR amplification using Red Taq DNA polymerase 2× Master mix (VWR, Oslo, Norway) according to the manufacturers protocol. The PCR reaction was carried out using 30 cycles, annealing temperature 58 °C (*AsChi18A, AsLPMO10A, AsLPMO10B*) or 56 °C (*VSAL_I0902*/*AsChi18Bp*) and 30 sec extension. To evaluate gDNA presence, samples prepared with no reverse transcriptase during cDNA synthesis (referred to as -RT control) was applied as template for primer pairs *AsLPMO10A* and VSAL_I0902.

PCR products were visualized by agarose gel electrophoresis of the total 20 µL PCR reaction in 1.3 % agarose 1xTAE electrophoresis buffer (Thermo scientific, Vilnius, Lithuania). The agarose was supplemented with peqGreen DNA/RNA dye (peqlab brand, VWR, Oslo, Norway) for visualization. After gel visualization, the gene expression was assessed as positive if the target gene was amplified in two out of three biological replicates and at the same time no amplification was observed in PCR samples prepared with the -RT controls. A complete list of primers used for amplification of target genes is shown in Table S4.

### Sample preparation and proteomic analysis

Biological triplicates of *Al. salmonicida* LFI1238 was incubated in 50 mL Asmm supplemented with 1 % β-chitin. At mid-exponential phase, cultures were harvested and fractioned into supernatant and pellet by centrifugation at 4 000 × *g* for 10 min at 4°C. β-chitin aliquots from the culture flasks were transferred to 2 mL Safe-Lock Eppendorf tubes (Eppendorf, Hamburg, Germany) and boiled directly for 5 min in 30 µL NuPAGE LDS sample buffer and NuPAGE sample reducing agent (Invitrogen™, CA, USA). Filter sterilized supernatant was concentrated using Vivaspin® 20 centrifugal concentrators (Vivaproducts, Littleton, MA, USA) by centrifugation at 4 000 rpm and 4 °C until it reached 1 mL concentrate. The bacterial pellet was lysed in 2 mL 1× BugBuster™ protein extraction reagent (Novagen), incubated by slow shaking for 20 min, followed by centrifugation and protein precipitation. Proteins were precipitated by adding trichloroacetic acid (TCA) to 10 % and incubation over-night at 4 °C. The precipitated proteins were harvested by centrifugation at 16 000 × *g* and 4°C for 15 min and washed twice in ice-cold 90 % acetone/0.01 M HCl. All final samples were boiled in 30 µL NuPAGE LDS sample buffer and sample reducing agent for 5 min and loaded on Mini-PROTEAN® TGX Stain-Free™ Gels (Bio-Rad laboratories, Hercules, CA, USA). Electrophoresis was performed at 300 V for 3 min using the BIO-RAD Mini-PROTEAN® Tetra System. Gels were stained with Coomassie Brilliant Blue R250 and 1×1 mm cube gel pieces were excised and transferred to 2 mL LoBind tubes containing 200 µL H_2_O. Sequentially, the gel pieces were washed 15 min in 200 µl H_2_O and decolored by incubating 2×15 min in 200 µL 50% acetonitrile, 25mM ammonium bicarbonate (AmBic). Next, reduction was performed by incubating the gel pieces in dithiothreitol (10 mM DTT/100mM AmBic) for 30 minutes at 56 °C and alkylation was done with iodo-acetamide (55 mM IAA/100mM AmBic) for 30 minutes at room temperature. After removal of the IAA solution, the gel pieces were dehydrated using 200µL 100% acetonitrile and digested using 30-45 µL of 10 ng/µL trypsin solution overnight at 37 °C. The next day, digestion was stopped by addition of 40 µL 1% trifluoroacetic acid (TFA). Peptides were extruded from the gel pieces by 15 minutes sonication and desalted using C18 ZipTips (Merch Millipore, Darmstadt, Germany), according to manufacturer’s instructions.

Peptides were essentially analyzed as previously described (95). In brief, peptides were loaded onto a nanoHPLC-MS/MS system (Dionex Ultimate 3000 UHPLC; Thermo Scientific) coupled to a Q-Exactive hybrid quadrupole orbitrap mass spectrometer (Thermo Scientific). Peptides were separated using an analytical column (Acclaim PepMap RSLC C18, 2 µm, 100 Å, 75 µm i.d. × 50 cm, nanoViper) with a 90-minutes gradient from 3.2 to 44 % [v/v] acetonitrile in 0.1% [v/v] formic acid) at flow rate 300 nL/min. The Q-Exactive mass spectrometer was operated in data-dependent mode acquiring one full scan (400-1500 m/z) at R=70000 followed by (up to) 10 dependent MS/MS scans at R=35000. The raw data were analyzed using MaxQuant version 1.6.3.3 and proteins were identified and quantified using the MaxLFQ algorithm (96). The data were searched against the UniProt *Al. salmonicida* proteome (UP000001730; 3513 sequences) supplemented with common contaminants such as human keratin and bovine serum albumin. In addition, reversed sequences of all protein entries were concatenated to the database to allow for estimation of false discovery rates. The tolerance levels used for matching to the database were 4.5 ppm for MS and 20 ppm for MS/MS. Trypsin/P was used as digestion enzyme and 2 missed cleavages were allowed. Carbamidomethylation of cysteine was set as fixed modification and protein N-terminal acetylation, oxidation of methionines and deamidation of asparagines and glutamines were allowed as variable modifications. All identifications were filtered in order to achieve a protein false discovery rate (FDR) of 1%. Perseus version 1.6.2.3 (97) were used for data analysis, and the quantitative values were log2-transformed, and grouped according to carbon source and condition (substrate/supernatant/pellet). Proteins were only considered detected if they were present in at least two replicates in at least one condition. All identified proteins were annotated with putative carbohydrate-active functions as predicted by dbCAN2 (98), biological functions (GO and Pfam) downloaded from UniProt, and for subcellular location using SignalP5.0 (99).

### Pseudogenes

Pseudogenes are gene sequences that have been mutated or disrupted into an inactive form over the course of evolution and is commonly thought of as “junk DNA”. The genome of *Al. salmonicida* LFI1238 contains a significant number of IS elements, and several genes are truncated and annotated as such pseudogenes. Since pseudogenes in general are believed to be non-functional, putative products of these are commonly not included in proteome databases. Consequently, a proteomic analysis towards the annotated proteome of *Al. salmonicida* LFI1238 will not detect products of these genes. To include these in our analysis, a few required steps were taken. Firstly, pseudogenes of three chitinases, a chitoporin and a chitodextrinase were selected as genes of interests based on the publication by Hjerde et al (29). Next, the truncated nucleotide sequence of a pseudogene was retrieved by searching the complete genome sequence annotation of *Al. salmonicida LFI1238* chromosome I (FM178379.1) for the specific gene locus. The gene locus of each selected pseudogene is shown in Table 7. The nucleotide sequences were translated to putative protein sequences using the translate tool at ExPASy Bioinformatics Resource Portal (100). The translate tool identifies potential start and stop codons of the query sequence by assessing reading frames 1-3 of forward and reverse DNA strand. Manually, putative peptides larger or equal to 6 amino acids were selected as supplement for the proteomic analysis. Pseudogene products of which unique peptides were identified was assigned a putative CAZy annotation using dbCAN2.

**Table 7.** Pseudogenes analyzed. . Gene locus, product name and CAZyme based name.

| Gene locus | Product | CAZyme based name |
| --- | --- | --- |
| *VSAL_I0763* | Chitinase A (fragment) | na |
| *VSAL_I0902* | Chitinase A (fragment) | *AsChi18Bp* |
| *VSAL_I1414* | Putative chitinase (pseudogene) | *AsChi19p* |
| *VSAL_I1942* | Chitinase (pseudogene) | *AsChi18Cp* |
| *VSAL_I2352* | Chitoporin (pseudogene) | na |
| *VSAL_I1108* | Chitodextrinase (fragment) | na |

## Data availability

The mass spectrometry proteomics data have been deposited to the ProteomeXchange Consortium via the PRIDE (101) partner repository with the dataset identifier PXD021397.

## ACKNOWLEDGEMENTS

This work was funded by grant 249865 (GV-K, AS, MØA and JSML) from the Norwegian Research Council and by a PhD fellowships from the Norwegian University of Life Sciences, Faculty of Chemistry Biotechnology and Food Science to GiM. BB has received the support of the EU in the framework of the Marie-Curie FP7 COFUND People Programme, through the award of an AgreenSkills fellowship (under grant agreement n° 267196). The authors would like to thank Simen Foyn Nørstebø (Faculty of Veterinary Medicine, Department of Food Safety and Infection Biology, Norwegian university of life sciences) for providing bacterial strains for deletion mutagenesis and for valuable advice regarding the protocol and Anne Cathrine Bunæs for assistance in purification of proteins.

## AUTHOR CONTRIBUTIONS

AS: Planned experiments, performed experiments, analyzed data, wrote the paper. JSML: planned experiments, wrote the paper. GiM: Planned experiments, performed experiments, analyzed data, wrote the paper. JSML: planned experiments, wrote the paper. SM: performed experiments, analyzed data, wrote the paper. BB: performed experiments, analyzed data, wrote the paper. MØA: performed experiments, analyzed data, wrote the paper. GeM: planned experiments, wrote the paper. GV-K: conceptualized the study, planned experiments, analyzed data, wrote the paper, supervised the study.

