## Supplementary material for "The fish pathogen *Aliivibrio salmonicida* LFI1238 can degrade and metabolize chitin despite major gene loss in the chitinolytic pathway"

### Supplementary figures

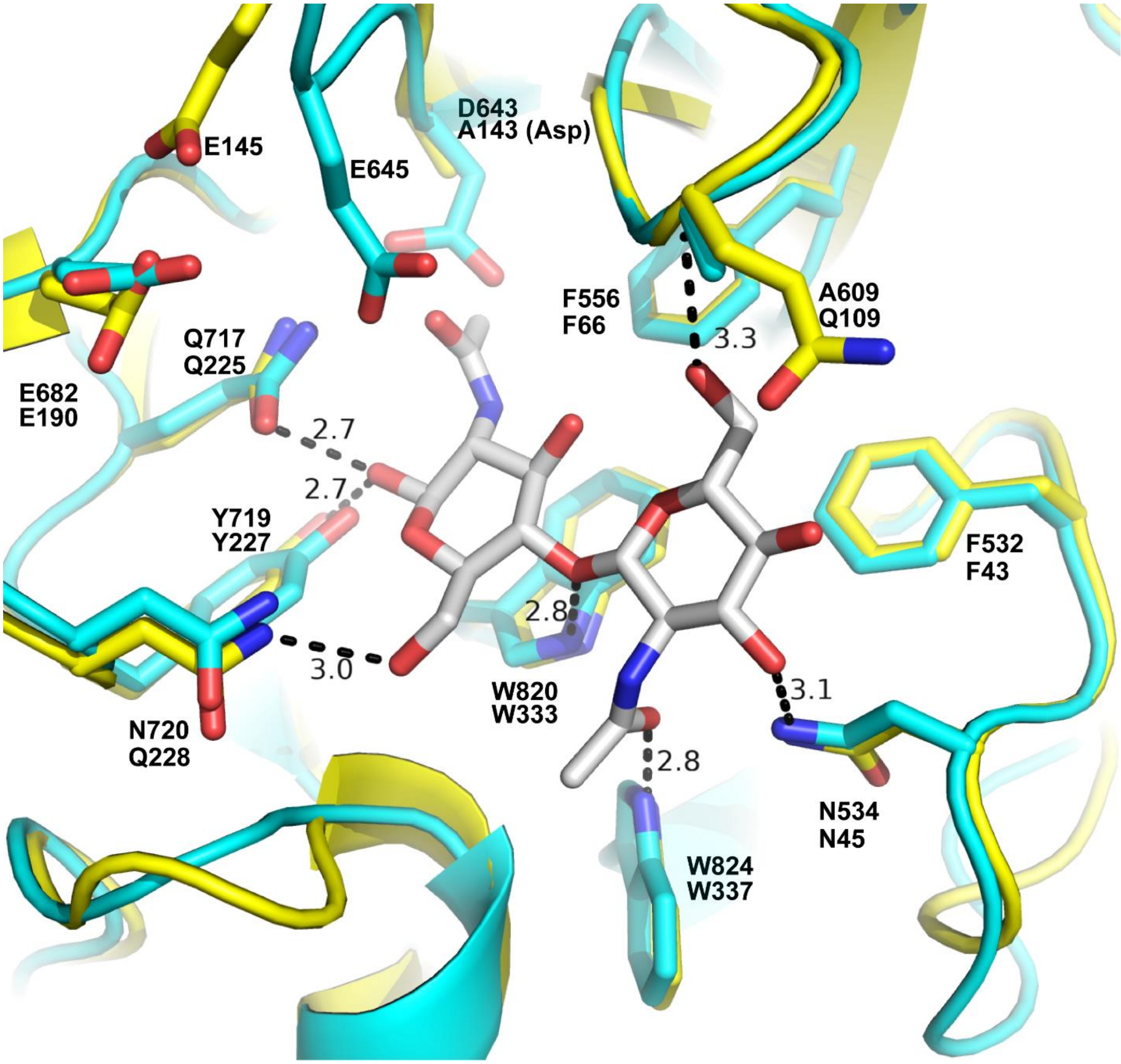


**Figure S1. Active site of the *As*Chi18A homology model superimposed on ChiNCTU2 (D143A variant).** Proteins are shown as cartoon representation with side chains shown in stick representation (*As*Chi18A colored cyan, ChiNCTU2 D143A [PDB id: 3N13] colored yellow). Side chains are labeled showing *As*Chi18A amino acid numbers above the ChiNCTU2 numbers. The chitobiose ligand bounbd in the -1 and -2 subsite of the ChiNCTU2 active is shown in stick representation with gray colored carbon atoms. Hydrogen bonds are illustrated by dashed lines and distances (Å) are indicated. It should be noted that the positioning of the ChiNCTU2 catalytic acid, E145, deviates from the position observed in the ChiNCTU2 wild type structure (which is more similar to the positioning of AsChi18A E645) due to the absence of an Asp at position 143.

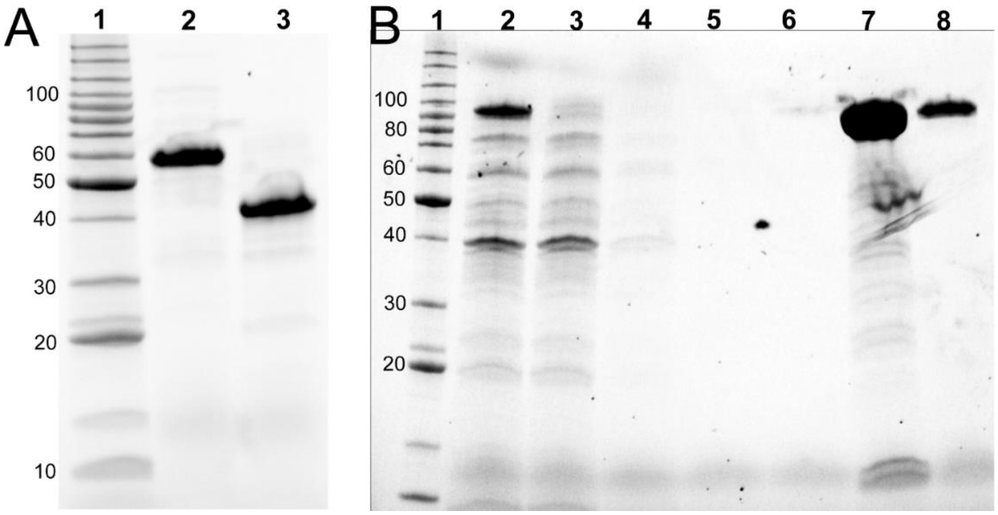


**Figure S2. Analysis of protein purity.** SDS-PAGE was used to determine protein purity. Panel A shows an SDS-PAGE gel with lanes displaying protein benchmark ladder (Invitrogen) in lane 1, *As*LPMO10A in lane 2 and *As*LPMO10B in lane 3. The SDS-PAGE gel in panel B displays the stages of protein purification for *As*Chi18A, showing the protein benchmark ladder in lane 1, cell free extract from an induced culture in lane 2, flow through in lane 3, wash fraction in lane 4-6 and the eluted protein in lanes 7 and 8. Only fractions containing highly pure protein were used in biochemical assays. All proteins used in biochemical assays were estimated to be >95% pure.

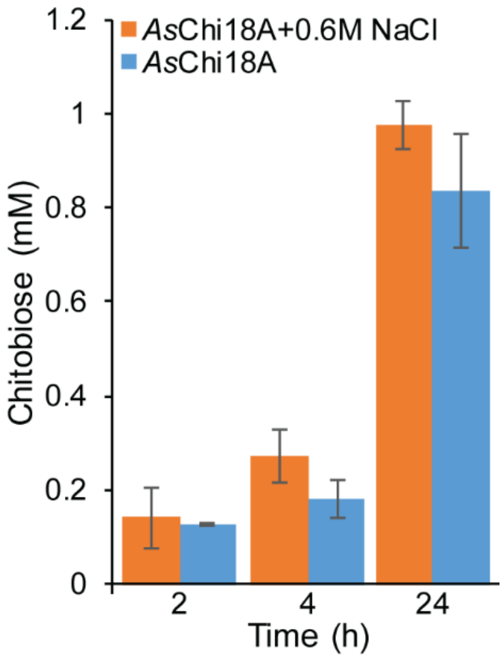


**Figure S3. Influence of NaCl on *As*Chi18A activity.** The amount of $(GlcNAc)_2$ (chitobiose) released from 10 mg/mL β-chitin in Tris-HCl pH 7.5 by 1.0 μM *As*Chi18A in the presence and absence of 0.6 M NaCl was evaluated at three time points (n=3). Reactions were incubated at 30°C.

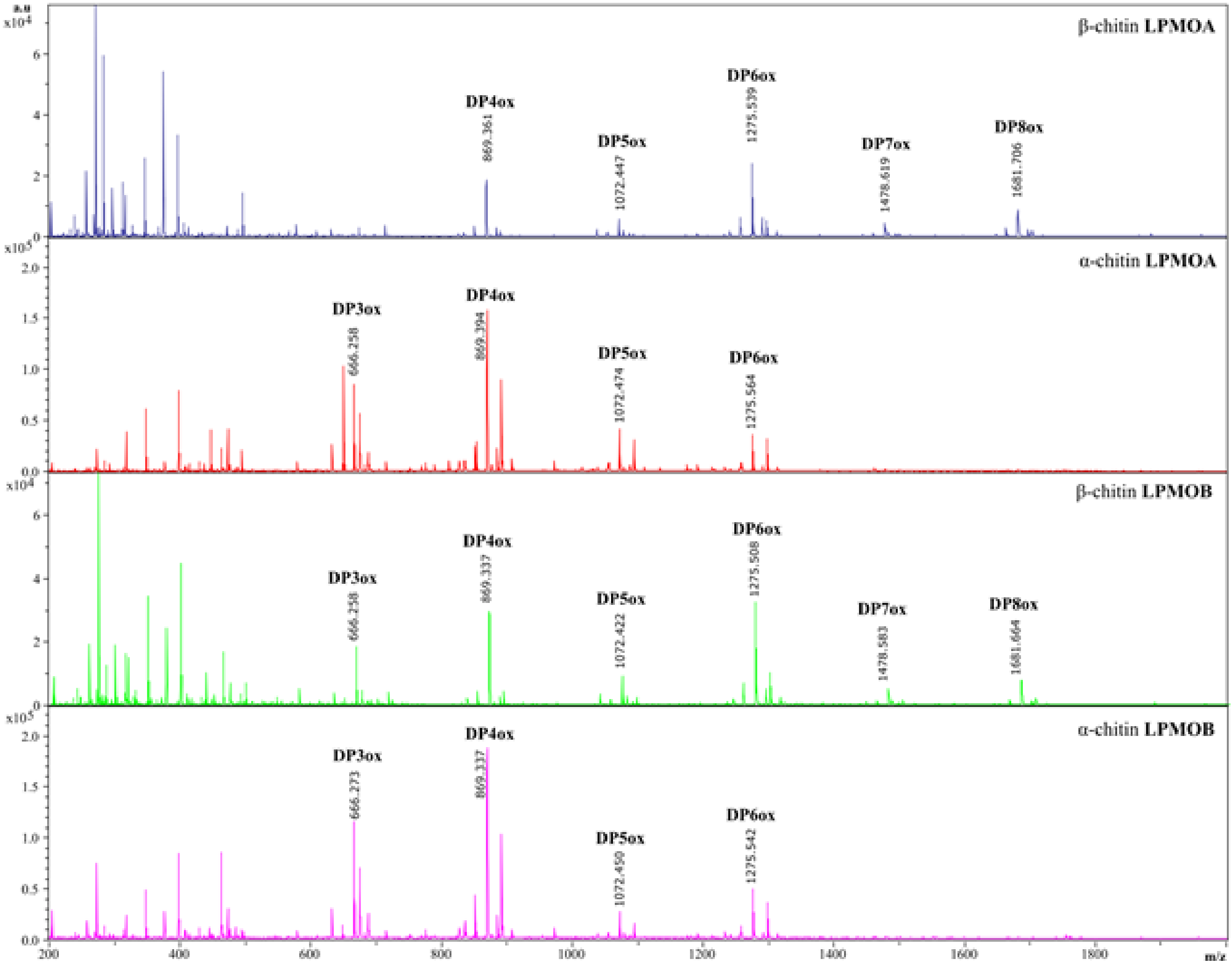


**Figure S4. MALDI-TOF MS analysis of oxidized products generated by *As*LPMO10A and -B from *A. salmonicida* on chitin (α and β).** The MS spectra show soluble C1 oxidized chito-oligosaccarides, i.e. aldonic acids. The degree of polymerization of each product is indicated by “DPn ox”, where n equals the number of monosaccharides in the chain. The main peaks are labelled with the respective masses.

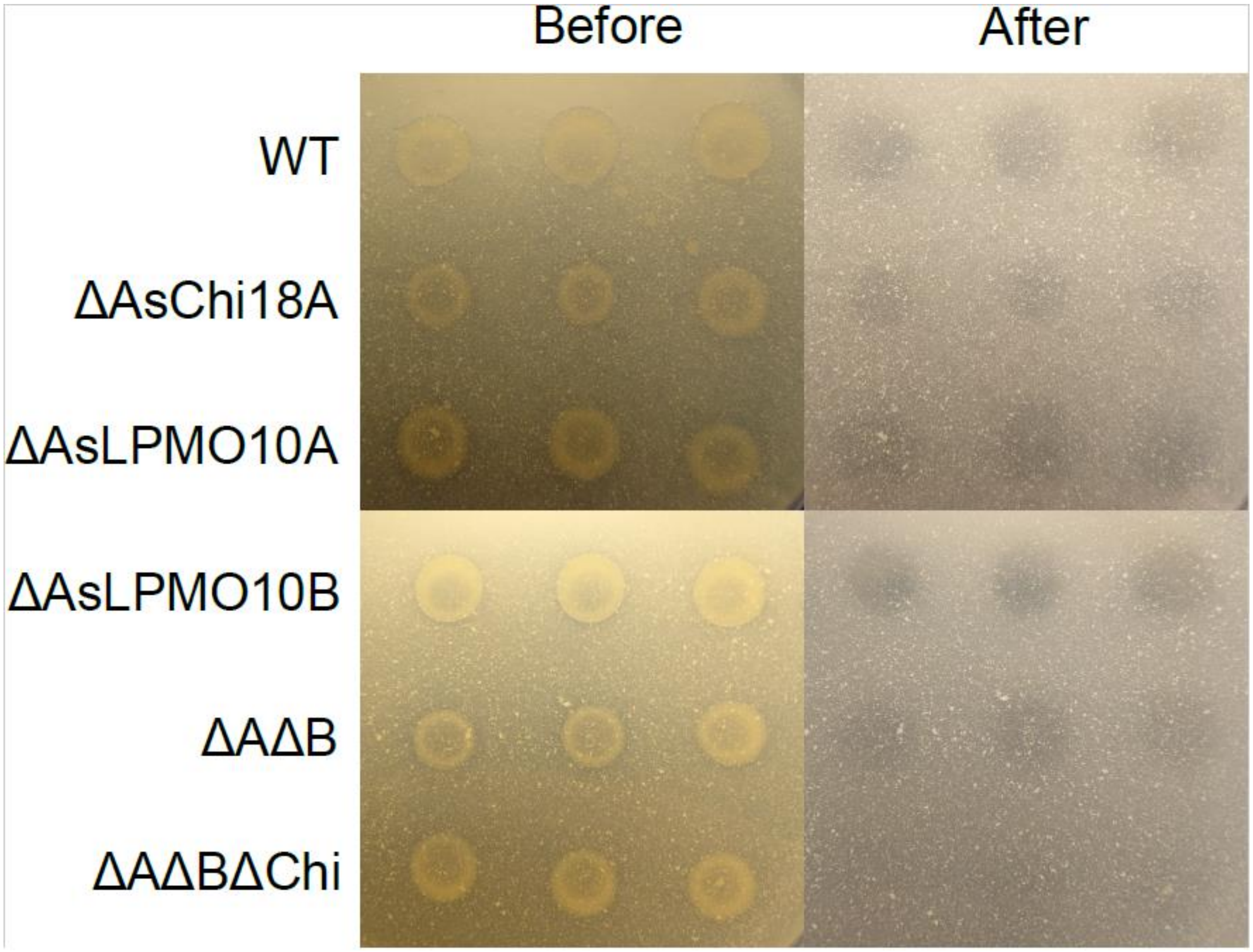


**Figure S5. Chitin degradation assay**. Images show photographs of agar plates containing LB25 supplemented with 2% colloidal chitin with *Al. salmonicida* variants (indicated on the left side of the image) spotted in triplicate and allowed to grow at 12 °C for 20 days. The photographs show the agar plates before (left) and after (right) the colonies had been removed by gentle washing. Halos indicate chitin degradation.

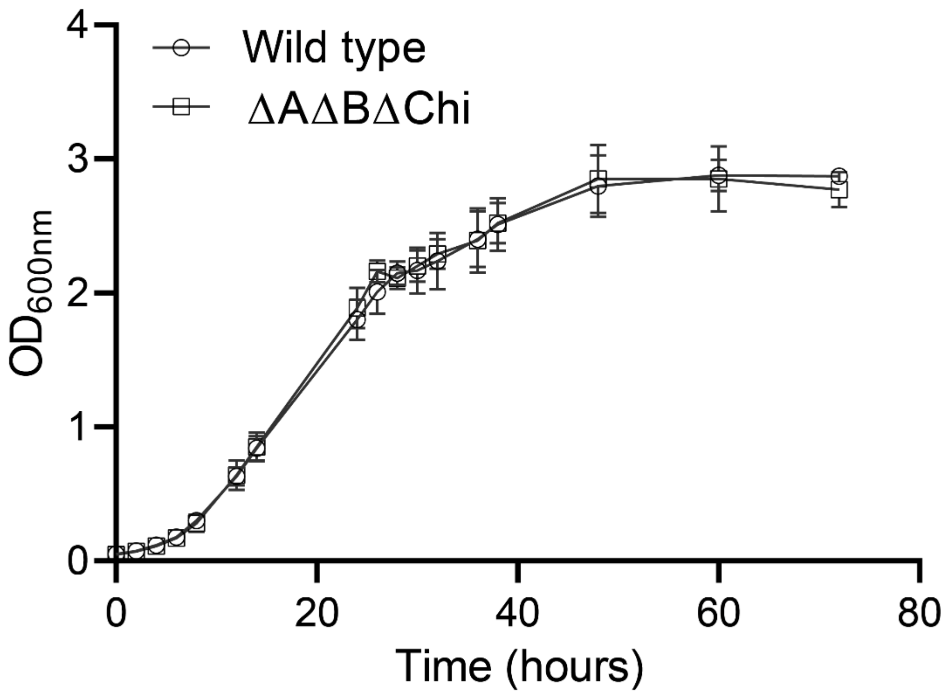


**Figure S6. Growth of *Al. salmonicida* LFI1238 variants.** Growth of the wild type *Al. salmonicida* LFI1238 strain compared to the triple knock-out strain (ΔAΔBΔChi) in LB25 broth. Standard deviation is indicated by error bars (n=3).

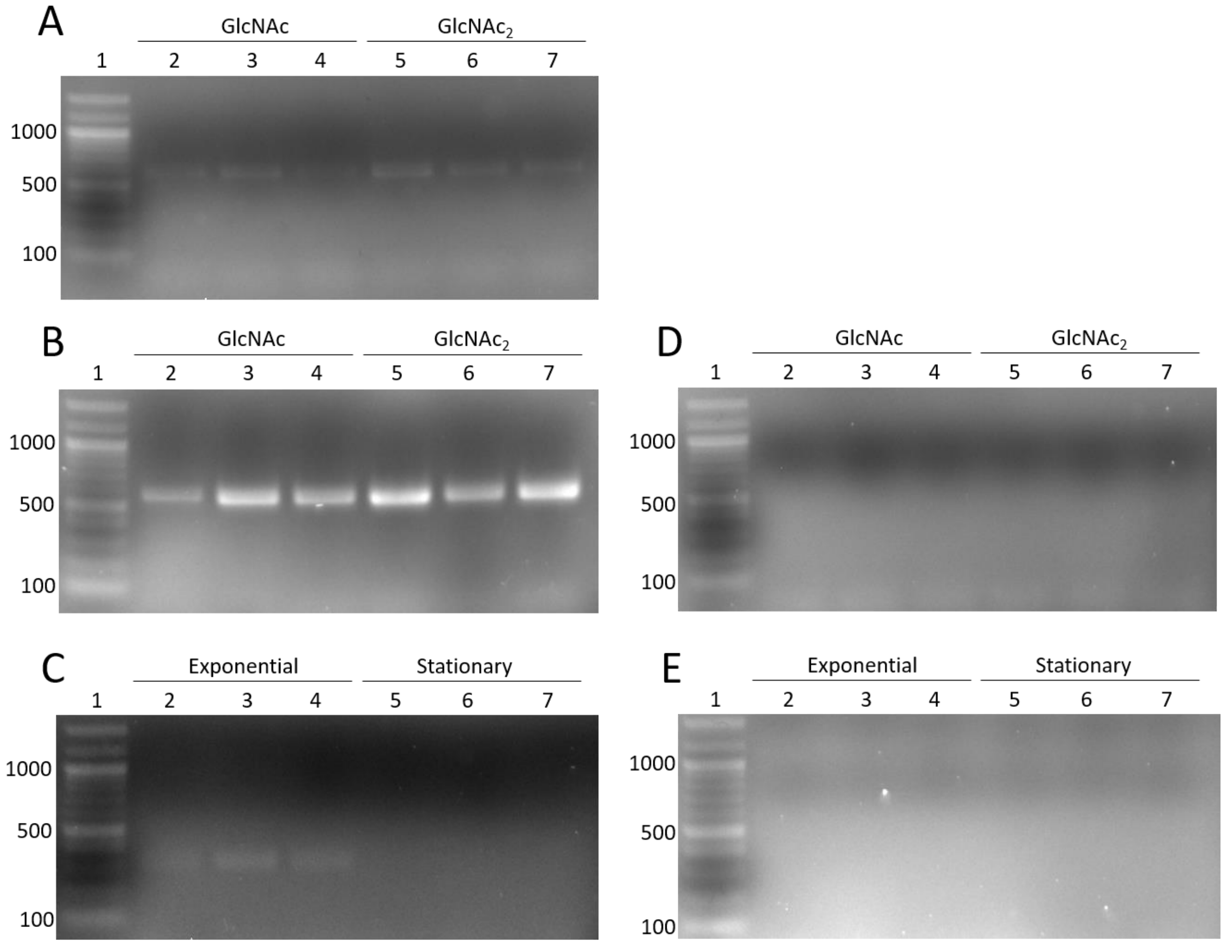


**Figure S7. Gene expression analysis by PCR amplification of cDNA.** Panel A and B shows the products formed in PCR experiments using cDNA from samples obtained during exponential growth in GlcNAc and GlcNAc$_2$ combined with primer pairs *GH18Expression* and *10AExpression*, respectively. Panel D shows the PCR experiments using -RT controls as template combined with primer pair *10AExpression.* The -RT templates used in Panel D corresponds to the cDNA applied to Panel A and B. Panel C shows the products formed in PCR experiments using cDNA from samples obtained during exponential and stationary growth in glucose combined with primer pair *I0902Expression*. In this case gene expression was evaluated as positive (+) during exponential growth (lane 2-4) and negative (-) during stationary growth (lane 5-7). Panel E shows the products formed in PCR experiments using the -RT samples corresponding to the cDNA template used in Panel C combined with primer pair *I0902Expression*. Lane 1; 100 bp DNA ladder, lanes 2-4 or 5-7; biological replicates within the same condition.

```
B6EGV7_ALISL     1 MKKTLLALTVSSIILSTPSLASAPNTNLNLMPYPQSVELKSGKLPIDNNFSIYIDGYKSE
D9ISD9_VIBHA     1 -----------------------------------MGGSGQGKITLDKSFSIYIKGYDSP

B6EGV7_ALISL    61 RIQQLAQRIIKRIEAQTGLPLLSPFSNSEKNATLIIRVNKAAKKEIQDNHIDESYQLSVN
D9ISD9_VIBHA    26 RVQFNAKRTMDRLYRQTGLPMLNWHAESEKDATLVIDIRNAPKSEVQDIHSDESYQLESR

B6EGV7_ALISL   121 QKQIILQAERPYGVIRGAETFLQLITTSKNGYSVPQIIIEDQPRFPWRGASFDSSRHFVS
D9ISD9_VIBHA    86 NGQIIIRSERPYGAFHGLETFLQLVTTDATGYFVPAVSIQDEPRFPWRGVSYDTSRHFIE

B6EGV7_ALISL   181 IETIKRQIDGFASAKMNVFHWHLWDDQAIRIQIESYPKLWQKTADGDVYTKEEIKDVIEY
D9ISD9_VIBHA   146 LDVILRQLDAMASAKMNVFHWHIWDDQAIRIQLDNYQSLWQDTADGDYYTKDEIRYVVNY

B6EGV7_ALISL   241 ARLRGIRVIPEISLPGHASGVAHAYPELMSGEGKQSYEQQRAWGVFVPLMNPLNPELYIF
D9ISD9_VIBHA   206 ARNLGIRVIPEISLPGHASAVAHAYPELMSGMGEQSYPHQRGWGVFEPLMDPTNPELYKM

B6EGV7_ALISL   301 FDNVFSEVTDLFPDEYIHIGGDEPNYQQWSNNKKIQAFIKENNIDGNRGLQSYLNARIEK
D9ISD9_VIBHA   266 LASVFDEVVELFPDEYFHIGGDEPNYQQWKDNPKIQQFIKDNNLDGERGLQSYLNTKVEQ

B6EGV7_ALISL   361 MLNDKGKKIMGWDEIWHKDLPTSIVIQSWRGHDSIGQAAKEGYAGLLSTGFYLDQPQPTS
D9ISD9_VIBHA   326 MLEQRGKKMTGWDEIWHKDLPTSIVIQSWQGHDSIGRAAKEGYQGILSTGYYLDQPQPTS

B6EGV7_ALISL   421 YHYRNDPMPKGPQVDDQQHQGESFETYTWQKPRGKGGPRKGTLTIITSKEGNARAFTDYN
D9ISD9_VIBHA   386 YHYRNDPMPKGITVDDQLHQGEKFATYDWVKPRNKGGPRTGNLTIIEATDGSYRAFTDYN

B6EGV7_ALISL   481 GKSRAKVDIIEYTKGESFRGHFDNFMSYTEFNLTMDNRQFSDGSYQLIGNVRWPTTGELT
D9ISD9_VIBHA   446 GKSREEVFIIEYVPGVKFKGHFDNFMSYTEFNYDFAGGKLKDSSYQLIGNVRWPATGELV

B6EGV7_ALISL   541 ASSSMKGTKIEKPTGGYPVALNEKEQVLILGGEAAIWAENYDDLTVEARIWPRTYAVGER
D9ISD9_VIBHA   506 ASSDMEGSVIPEPNGGYPAELTEKEQPLILGGEITIWGENLDSMTIEQRLWPRSYAVAER

B6EGV7_ALISL   601 LWSAESLTDEDSMYKRLEVMNNWATISVGLQHQANSYKQYLRLA---------SFANYAE
D9ISD9_VIBHA   566 LWSSQDLTDERSMYKRMKVMDTWSEISLGLRHHADANIMLKRLANGADETPLQTLAKYIE

B6EGV7_ALISL   652 PAQYYARNWAKFNATDPKGELYNQYERLNRFVDAVPVESLAVLKMMSL-ATTATNTPSDL
D9ISD9_VIBHA   626 PAQYYARHWEKWISTPNEGDLYNQYERLNRFADALPVESIAVYEMQDLVAAYALGDKTAL

B6EGV7_ALISL   711 QELKRHYQTVLDSAIKSKPIFENNIASIDTAPLAEKHKEVATAALELIAILEKGNKPTST
D9ISD9_VIBHA   686 DALAMHYQSVKLAAQQAKPIFAANVASVETVPVAEAAIKVADLGLTLIELAKQGSDISQS

B6EGV7_ALISL                771         QLRHVISVTTTASGMYDEMIVAIVRPTEELVRQLRKQ
D9ISD9_VIBHA   746 DAEAYQRIINENAIIFDETIVAIVVPTEQLLHTLTD-
```

**Figure S8.** Pairwise sequence alignment of the *Al. salmonicida* GH20 (UniProt ID B6EGV7) and *Vibrio harvey* GH20 *Vh*NAG1 (UniProt ID D9ISD9). The alignment was made using the EMBOSS pairwise sequence alignment tool using default parameters. The Alignment was formatted using BoxShade. Based on data from W. Suginta et al. (1), catalytic amino acids are shown in red shading and amino acids involved in substrate binding are shown in blue shading.

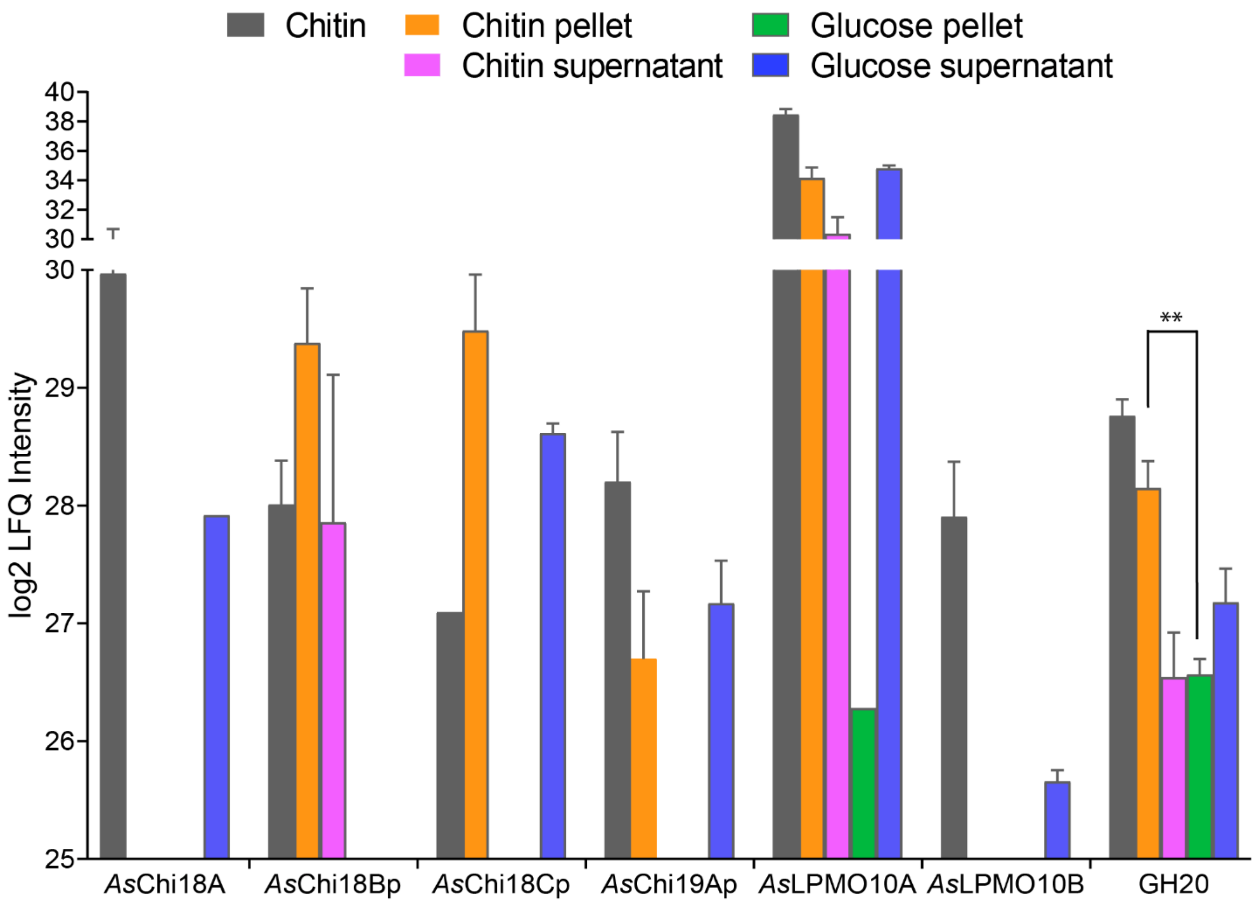


**Figure S9**. ***Al. salmonicida* protein abundances.** The abundance of selected *Al. salmonicida* proteins related to chitin catabolism displayed in bar chart format as a supplement to Figure 9. Log2 LFQ values shown represent the average values obtained from the label free proteomics data (Supplementary Dataset 1). Error bars are shown for proteins that were detected in two or three biological replicates. The GH20 β-*N*-acetylhexosaminidase was calculated to have a statistically significant 1.58 log2 fold higher abundance during growth on chitin compared to glucose ($p=0.0082$; paired two-tailed t test).

### Supplementary tables

**Table S1.** Growth rate measurements and max cell density of *Al. salmonicida* cultivated in glucose, GlcNAc and $GlcNAc_2$

| Carbon source | Rate constant μ (hours$^{-1}$) | Generation time (hours) | Max cell density ($OD_{600}$) |
|---|---|---|---|
| Glucose | 0.065 ± 0.025 | 5.36 ± 2.17 | 2.63 ± 0.094 |
| GlcNAc | 0.069 ± 0.029 | 5.16 ± 2.00 | 1.31 ± 0.022 |
| $GlcNAc_2$ | 0.055 ± 0.021 | 4.95 ± 0.89 | 1.58 ± 0.145 |

Mean ± SD of three biological replicates.

**Table S2.** Identified CAZymes sorted by their putative biological processes, according to gene ontology (GO) annotations.

| CAZy | Uniprot | Biological process |
|---|---|---|
| | | **Carbohydrate metabolic process [GO:0005975]** |
| AA10 | B6EQJ6 | *As*LPMO10B |
| CBM73;GH18 | B6EH15 | *As*Chi18A |
| GH13_19 | B6EGT4 | Alpha-amylase |
| GH20 | B6EGV7 | Putative beta-N-acetylhexosaminidase |
| GT35 | B6EQ29 | Alpha-1,4 glucan phosphorylase |
| GH77 | B6EQ30 | 4-alpha-glucanotransferase |
| GH3 | B6ERJ6 | Beta-hexosaminidase* |
| | | **Chitin binding [GO:0008061]** |
| AA10;CBM73 | B6EQB6 | *As*LPMO10A |
| | | **Cell cycle, cell division, protein import [GO:0007049/ 0051301/ 0017038]** |
| PL22 | B6EGK3 | Tol-Pal system protein TolB |
| | | **Formaldehyde catabolic process [GO:0046294]** |
| CE1 | B6EH03 | S-formylglutathione hydrolase |
| | | **Glycogen biosynthetic process [GO:0005978]** |
| GT5 | B6EQL7 | Glycogen synthase |
| | | **Lipid A biosynthetic process [GO:0009245]** |
| CE11 | B6ELH0 | UDP-3-O-acyl-N-acetylglucosamine deacetylase |
| GT19 | B6EJW7 | Lipid-A-disaccharide synthase |
| | | **Lipopolysaccharide biosynthetic process [GO:0009103]** |
| GT9 | B6EPB8 | ADP-heptose-LPS heptosyltransferase II |
| | | **Peptidoglycan metabolic process [GO:0000270]** |
| GH23 | B6EJV5 | Membrane-bound lytic murein transglycosylase D |
| GH23 | B6EGC8 | Soluble lytic murein transglycosylase |
| | | **Trehalose catabolic process [GO:0005993]** |
| GH13_29 | B6ERJ9 | Trehalose-6-phosphate hydrolase |
| | | **Not assigned** |
| GH103 | B6EIW0 | Putative exported protein |
| GT2 | B6EKR9 | Putative glycosyl transferase |
| GT51 | B6EM36 | Penicillin-binding protein 1A |

*Also, Cell cycle [GO:0007049];cell division [GO:0051301];cell wall organization [GO:0071555];peptidoglycan biosynthetic process [GO:0009252];peptidoglycan turnover [GO:0009254];regulation of cell shape [GO:0008360]

**Table S3.** Primers designed for construction of flanking regions and fusion product, sequencing and selection/verification of transconjugates and mutants.

| Primer | Sequence 5'-3' |
|---|---|
| *AsGH18*_For | GCTGATGGCGTGATCAAC |
| *AsGH18*_Rev | GGCGCGTGCTAATTTCAA |
| *AsLPMO10A*_For | GGCTGCTATTGTCACAGAATA |
| *AsLPMO10A*_Rev | AAGCCTAATAAAGCACACCCA |
| *AsLPMO10B*_For | GATGAGGTGTACCATCTTGAA |
| *AsLPMO10B* _Rev | TGTAATAGAATGTCACCAGCA |
| pDM4_Seq_F | CGGGAGAGCTCAGGTTAC |
| pDM4_Seq_R | GGCTTCTGTTTCTATCAGCT |

**Table S4.** Primers applied for amplification of target genes using cDNA.

| *Primer* | *Sequence (5'- 3')* | *Product size (bp)* |
|---|---|---|
| *GH18Expression_F* | AGTCAAGCATCAGCCAAGAAAG | 566 |
| *GH18Expression_R* | TAAGGCAAGGCTCGATCCAG | |
| *10AExpression_F* | ATTCGGTCCTGCTGATGG | 565 |
| *10AExpression_R* | ATTTGCTTGACCTTGTGTTGC | |
| *10BExpression_F* | TCAAGCGTGTCAGTCTGC | 441 |
| *10BExpression_R* | TGCCAACGAGTGTAGAGC | |
| *I0902Expression_F* | ATGCACAAGGTCGATCTG | 297 |
| *I0902Expression_R* | ATGGGATGTACTTGTCGC | |
